# Virtual phenotypic screening discovers novel scaffolds inhibiting the PI3K/mTOR pathway

**DOI:** 10.64898/2026.06.10.731476

**Authors:** Alexander P. Wu, Heming Yao, Burkhard Hoeckendorf, Garrett Gaskins, Nont Kosaisawe, Ziqing Liu, Phil Hanslovsky, Oleg Mayba, Nicholas Skelton, Gabriele Scalia, John G. Moffat, Tommaso Biancalani, Jan-Christian Hütter, David Richmond

**Author notes:** These authors contributed equally to this work (Alexander P. Wu, Heming Yao, Burkhard Hoeckendorf, Garrett Gaskins). Work performed while employed at Genentech (Tommaso Biancalani, Jan-Christian Hütter).

## Abstract

Phenotypic drug discovery has yielded many first-in-class small-molecule drugs by discovering modulators of disease phenotypes in physiologically relevant cellular systems. However, high-content phenotypic assays lack the ultra-high-throughput scalability of target-based screens. Recent advances in virtual screening present an opportunity to address this bottleneck, but have been limited to simple phenotypes like viability, restricted to small repurposing libraries, or lack in-depth biological validation. Here, we present PhenoCompass, a multimodal co-embedding model that aligns compound structures and high-content phenotypic imaging to enable virtual phenotypic screening over billion-compound libraries. Following training on the Joint Undertaking in Morphology dataset with more than 100,000 Cell Painting compound profiles, retrospective validation with historical biochemical high-throughput screening data demonstrates that PhenoCompass ranks compounds according to their biochemical target engagement. Leveraging PhenoCompass, we performed a prospective screen of 3.8 billion Enamine REAL compounds for inhibitors of PI3K/mTOR pathway, a critical signaling cascade whose aberrant activation is a common tumor driver. This search identified 11 novel compounds with pathway-consistent Cell Painting readout and diverse scaffolds, a 54-fold enrichment over the training set. Orthogonal validation experiments using a FOXO3A reporter assay and direct kinase inhibition confirmed seven structurally novel inhibitors with distinct mechanisms of action. These results highlight the convergence of diverse molecular target profiles onto a shared morphological pathway signature and establish PhenoCompass as a robust framework for high-content phenotypic virtual screening.

## Introduction

A fundamental challenge in drug discovery is the profound disparity between the scale of drug-like chemical space, estimated to exceed 10^60^, and the throughput of physical screening. Large-scale physical hit-finding methods range from arrayed screens (typically 10^5^ – 10^6^ compounds), including cell-based phenotypic and target-based Ultra-High-Throughput Screens (uHTS), to pooled DNA-encoded library target-binding selection (up to 10^12^ library members). These modalities present trade-offs between scale and physiological relevance that impact the risk and cost of progressing unoptimized hits to efficacious clinical candidates. In particular, phenotypic drug discovery (PDD) has been a major historical driver of first-in-class small-molecule drugs, complementing target-based drug discovery (TDD) through its use of unbiased functional readouts, allowing it to identify drugs acting through novel mechanisms, complex cellular pathways, and even polypharmacology^1–4^. However, PDD is the least scalable approach, requiring trade-offs between the biological complexity and translational validity of assays with the scale of libraries that can be screened^5^.

In TDD, virtual screening has demonstrated the potential to overcome scale limitations, primarily through structure-based docking methods, enabling computational prioritization of candidate molecules from vast *in silico* chemical libraries^6^. By contrast, the absence of a pre-specified target hypothesis in PDD makes purely structure-based methods infeasible. Nevertheless, ligand-based machine learning methods have recently shown promise for predicting simple phenotypes, such as bacterial viability, directly from chemical structure for virtual screening of antibiotics^7,8^. Extending this paradigm to specific disease states in mammalian cells has tremendous potential, but will require rich, high-dimensional phenotypic descriptors capable of capturing pathway-level biology.

In this context, Cell Painting has emerged as a cost-effective and broadly generalizable assay for large-scale, high-content phenotypic screening^9^. Cell Painting captures images of cellular organelles in a hypothesis-free manner, providing a high-content morphological “fingerprint” of biological perturbagens, and can distinguish phenotypic effects across compounds with diverse mechanisms of action (MoAs)^10–13^. Recent studies combining large-scale Cell Painting datasets with multimodal contrastive learning have successfully co-embedded chemical structures with their corresponding phenotypic responses, enabling bidirectional retrieval between structures and phenotypes^14–17^. Such methods suggest the feasibility of predicting phenotypic responses directly from chemical structures; however, they have yet to demonstrate prospective virtual screening of multi-billion compound libraries or pathway-specific discovery.

Here, we present PhenoCompass, a multimodal deep learning framework designed to enable virtual phenotypic screening at the billion-compound scale (**Fig. 1a**). Leveraging Geometric Multimodal Contrastive (GMC) representation learning on the Joint Undertaking in Morphological Profiling (JUMP) dataset^18^ (>100,000 compound-phenotype pairs profiled in U2OS osteosarcoma cells), PhenoCompass constructs a joint embedding of chemical topology and cellular morphology. We retrospectively validated PhenoCompass on historical HTS data, recovering potent inhibitors of HDAC2, JAK3, and PIK3CA in an affinity-dependent manner. We then performed a prospective virtual screen of the Enamine REAL library (3.8 billion compounds) for modulators of the PI3K/mTOR pathway, a signaling cascade frequently dysregulated in cancer^19–22^. From 396 prioritized candidates, 11 elicited PI3K/mTOR-consistent activity by Cell Painting, a 54-fold enrichment relative to the hit rate in the JUMP training set. Orthogonal validation with a FOXO3A reporter assay and kinase inhibition assays confirmed seven novel and structurally diverse PI3K/mTOR inhibitors with distinct mechanisms of action. Importantly, strong concordance across validation assays establishes PhenoCompass as a versatile framework for virtual phenotypic screening at unprecedented scale, effectively bridging the gap between compound diversity and phenotypic discovery.

**Fig. 1:**
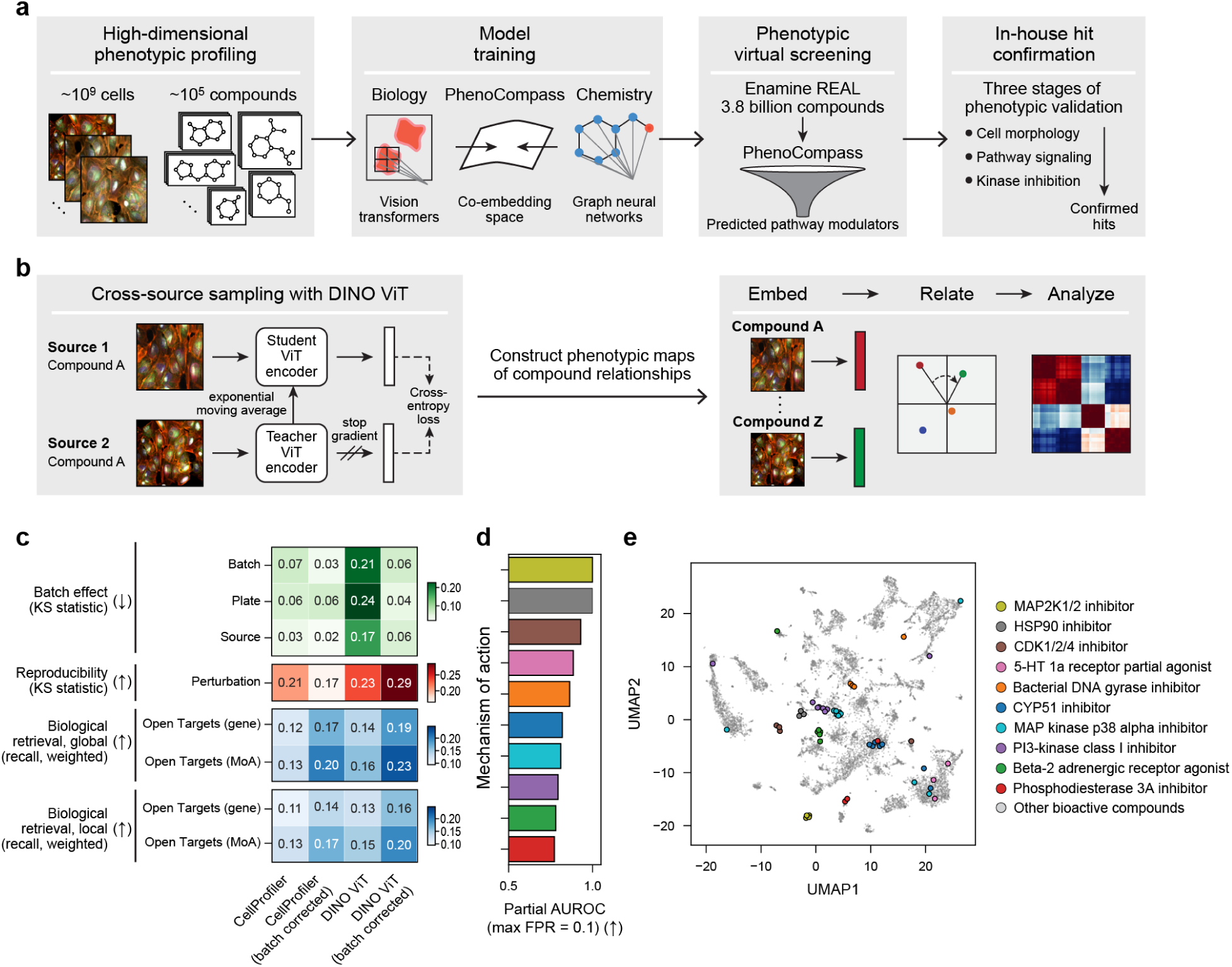
PhenoCompass links chemical compounds to biological effects through morphological phenotypic maps. **a,** PhenoCompass virtual screening pipeline for discovering pathway-specific chemical modulators. From left: A large training corpus of compound perturbations and associated phenotypic profiles is used for model training. PhenoCompass co-embeds vision transformer representations of biology with graph neural network representations of chemistry. This trained model enables phenotypic virtual screening on large-scale make-to-order libraries for pathway modulators, which are then validated in three stages of additional profiling. **b,** Pipeline for constructing and evaluating phenotypic maps from JUMP Cell Painting images. From left: Image representation learning leverages cross-source sampling using the DIstillation with NO Labels (DINO) architecture. The resulting representations are then used to embed compounds and analyze their phenotypic similarity via cosine similarity. **c,** Batch-corrected DINO map provides the highest biological recall and reproducibility while minimizing batch effects. Different evaluation metrics (y axis, **Methods**) across different phenotypic map variants (x axis), colored by the value of the metric (color bar). **d,e,** DINO phenotypic map groups compounds by MoA. **d,** Top 10 mechanisms of action (MoAs) in the JUMP phenotypic map (y axis, colors correspond to legend in (e) ranked by partial area under the receiver operating characteristic curve (pAUROC) with a maximum false positive rate (FPR) of 0.1 (d, x axis bar plot). **e,** Uniform Manifold Approximation and Projection (UMAP, scatter plot) of mean-aggregated compound phenotypic profiles (dots). Colored dots correspond to top-10 MoAs (legend); light gray dots are sampled from a background of bioactive compounds with strong effects (**Methods**). Arrows: higher (↑) or lower (↓) values represent better performance.

## Results

### A phenotypic map of JUMP resolves diverse compound mechanisms of action

To accurately link chemical structures with phenotypic responses, it is first necessary to construct a robust response map that captures biological heterogeneity while mitigating batch effects. To this end, and to fully leverage the unprecedented scale of the JUMP Cell Painting dataset^18^ despite its multi-center batch effects^23^, we constructed a high-fidelity phenotypic map using DINO, a self-supervised method for image representation learning with vision transformers^24–26^. We addressed batch effects through cross-batch sampling^27–29^ (**Fig. 1b**), followed by feature normalization, dimensionality reduction, and linear batch correction^30^ (**Methods**). Rigorous quality control revealed that shared well coordinates across replicate plates severely confounded the genetic perturbation data, artificially inflating their reproducibility rates (>90%) relative to previously reported rates (∼35%)^30^, with particularly high rates (99%) for genes unexpressed by U2OS cells (**table S1, Methods**). Consequently, we restricted our analysis entirely to compound perturbations and applied stringent filtering to eliminate high-variance sources and low-cell-count wells (**fig. S1, Methods)**. By computing pairwise cosine similarities between corrected representations of these compound perturbations, we established a high-fidelity phenotypic map encompassing 613,676 Cell Painting wells and representing 108,836 distinct compounds across eight sources.

To evaluate these representations, we benchmarked our DINO-derived maps against standard CellProfiler^31^ features across three criteria: batch effect reduction, replicate reproducibility, and the recall of known compound targets and mechanisms of action (MoAs)^23,30^. Consistent with previous findings that deep representation learning methods outperform engineered feature extraction^25,32–34^, DINO combined with a batch correction pipeline significantly outperformed all CellProfiler baselines, achieving a >10% improvement in both biological recall and reproducibility metrics (**Fig. 1c**). In the resulting JUMP phenotypic map, compounds with shared MoAs clustered robustly, capturing known strong-phenotype classes with high precision, such as PI3K inhibitors, MEK inhibitors, beta-2 adrenergic receptor agonists, and cytochrome P450 inhibitors (partial AUROC ≥ 0.78, max FPR = 0.1; **Fig. 1d**)^25,35–37^. Although visualization confirmed distinct MoA-specific clustering, residual heterogeneity remained, likely reflecting distinct off-target pharmacological effects or unresolved technical artifacts^23^ (**Fig. 1e**, **fig. S2**).

### PhenoCompass accurately predicts morphological response to compound perturbations

We framed the challenge of predicting cellular phenotypic responses from chemical structures as a multimodal alignment task^14–17,38^ and developed PhenoCompass, a self-supervised model that aligns information from molecular structures and Cell Painting images using geometric multimodal contrastive (GMC) learning^39^ (**Fig. 2a**). Unlike standard contrastive approaches (e.g., CLIP)^40,41^, GMC employs a joint-modality encoder as an anchor to align modality-specific representations with a fused representation rather than directly with each other, preserving both chemical and biological information (**Methods**, **fig. S3**). To rigorously evaluate generalization to structurally novel compounds, or “scaffold hopping”^42^, we evaluated PhenoCompass using scaffold-cluster splits^8^. By holding out entire groups of compounds with structurally similar scaffolds, rather than individual compounds with identical scaffolds like in standard scaffold splits, we generated highly out-of-distribution test sets, ensuring our benchmarks reflect real-world discovery challenges (*p* < 10^−92^, one-sided Wilcoxon signed-rank test between distributions of maximum Tanimoto similarity (TS) to train set, **Methods, fig. S4**).

**Fig. 2:**
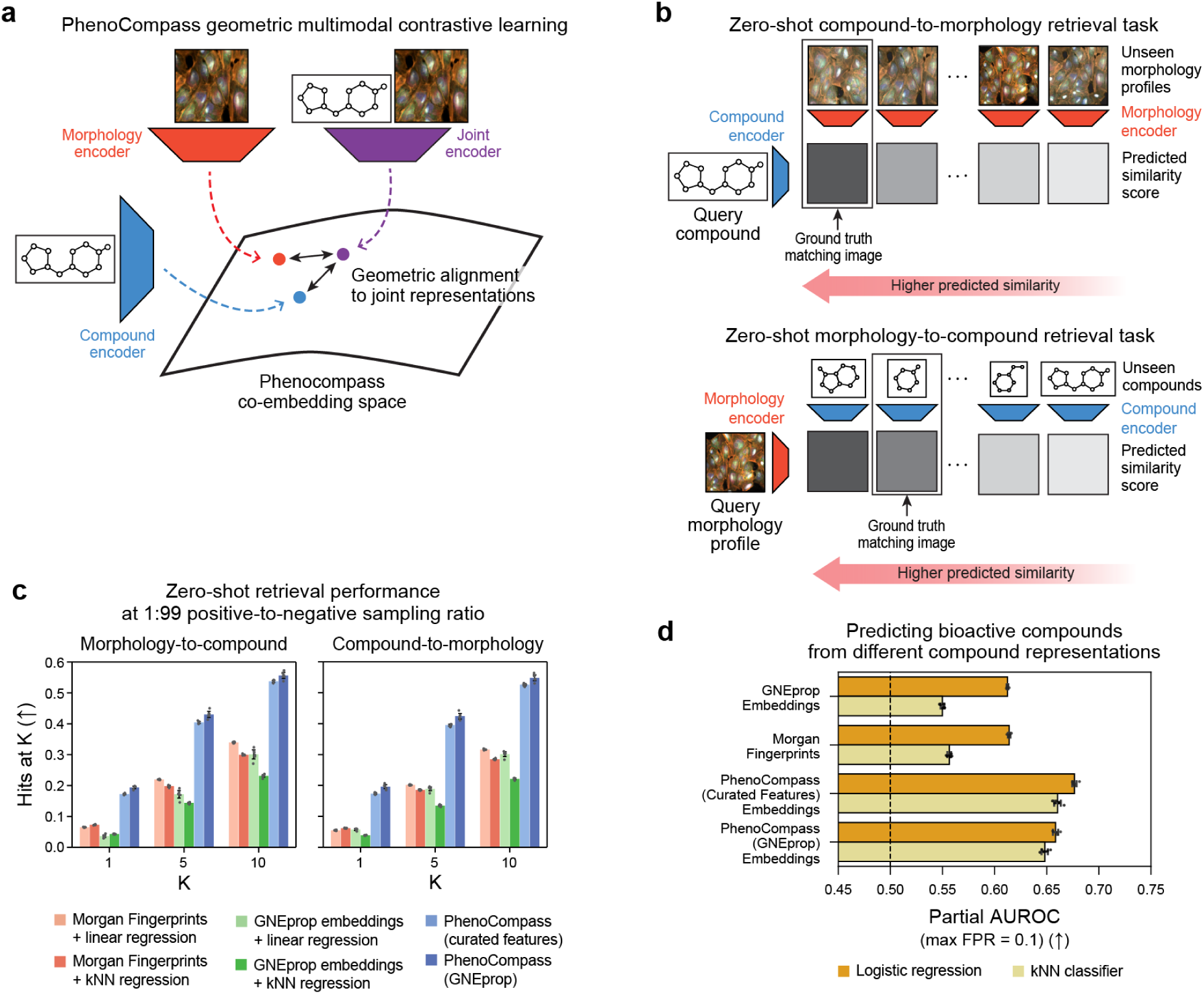
Geometric multimodal learning enables phenotypic predictions for structurally novel compounds. **a**, Schematic of the geometric multimodal contrastive (GMC) learning framework for multimodal representation learning with PhenoCompass. Morphology and compound encoders embed images and chemical structures, respectively. A joint encoder embeds both modalities, and the modality-specific representations are geometrically aligned to these joint representations within the PhenoCompass co-embedding space. **b**, Schematic of the zero-shot compound-to-morphology and morphology-to-compound retrieval tasks. Top: Given a query compound, its structure is encoded and compared against unseen morphology profiles to predict the correct matching image. Bottom: Given a query morphology profile, the image is encoded and compared against unseen compounds to predict the correct matching compound. **c, d**, PhenoCompass outperforms baseline representations on zero-shot retrieval and bioactivity prediction tasks. c, Assessment of PhenoCompass and baseline models for the morphology-to-compound (left) and compound-to-morphology (right) retrieval tasks across six scaffold-cluster train-test splits at a 1:99 sampling ratio (**Methods**). Retrieval performance (y axis) is evaluated as the proportion of correctly matched compound-image pairs among the top K predictions (Hits at K) for K = 1, 5, and 10 (x axis) across varying models (bar colors, legend). d, Assessment of logistic regression and k-nearest neighbors (kNN) classifier models for predicting bioactive compounds using GNEprop embeddings, Morgan Fingerprints, and PhenoCompass compound representations. Prediction performance (x axis) is evaluated using partial area under the receiver operating characteristic curve (pAUROC, max FPR = 0.1) for different representation methods (y axis) grouped by classifier type (bar colors). Bars: mean ± 2 s.e. of six splits. Arrows: higher (↑) or lower (↓) values represent better performance.

We first evaluated PhenoCompass’s capacity to map novel chemical spaces to phenotypic effects via zero-shot cross-modal retrieval. For each scaffold-cluster split, we assessed whether a true compound-image pair could be accurately identified against 99 decoys (**Fig. 2b**). We compared two distinct compound encoders: PhenoCompass (Curated Features), utilizing Morgan, Avalon, ErG, and RDKit descriptors^7,43,44^, and PhenoCompass (GNEprop), an end-to-end trained graph neural network leveraging the GNEprop architecture^8^. Each PhenoCompass model was also compared against four baseline models: linear and k-nearest neighbors (kNN) regression, each trained on Morgan Fingerprints or pre-trained GNEprop features as alternative compound representations (**Methods**). PhenoCompass models consistently outperformed all baselines in both compound-to-morphology and morphology-to-compound retrieval (**Fig. 2c**, **Methods**). Specifically, PhenoCompass (GNEprop) achieved top-1, top-5, and top-10 recall accuracies of 19%, 42%, and 55%, respectively. This significantly exceeded the random baselines (1%, 5%, 10%) and demonstrated up to a 3.2-fold improvement in hit rates over the next-best baseline, while remaining robust to plate-level batch effects (**Methods, fig. S5a**).

Motivated by these retrieval capabilities and the sensitivity of Cell Painting as a bioactivity readout^45,46^, we next evaluated the utility of PhenoCompass representations for predicting the bioactivity of unseen compounds (**Methods**). Using the same test set based on scaffold-cluster splitting, we compared supervised logistic regression probes and kNN classifiers trained on compound representations from both PhenoCompass variants as well as Morgan Fingerprint and pre-trained GNEprop baselines. We evaluated performance using partial AUROC (pAUROC; max FPR=0.1) to prioritize the top-ranked predictions for virtual screening. Logistic regression models trained on PhenoCompass representations yielded significantly higher pAUROC values than those trained on Morgan Fingerprints or pre-trained GNEprop embeddings (*p* < 0. 05, paired one-sided Wilcoxon rank-sum test, **Fig. 2d**). Notably, PhenoCompass (Curated Features) achieved the highest average pAUROC of 0.68, compared to 0.61 for the best baseline models. The kNN classifiers consistently underperformed the linear probes, indicating that the native geometric spaces of standard representations are less informative for bioactivity than the targeted subspaces identified by supervised models.

### PhenoCompass accurately predicts pathway modulators from compound structure with few-shot learning

Aberrant signaling pathways frequently drive disease pathogenesis, making the discovery of pathway-specific chemical modulators a critical drug discovery objective^1,47^. To identify structurally novel pathway modulators, we deployed PhenoCompass in a few-shot virtual screening framework. By leveraging known pathway modulators as “anchors” and embedding them via PhenoCompass’ joint-modality encoder, we can query novel libraries using only the candidate compounds’ structural representations, where higher cosine similarity to the anchor embeddings indicates greater predicted similarity to the desired phenotype (**Fig. 3a**). Anchor sets were generated by this approach for six diverse target classes: PI3K/mTOR, JAK/ROCK, CDK, HDAC, MAPK, and HSP90. Each set comprised 7 to 17 structurally distinct compounds (intra-set Tanimoto similarities ranging from 0.03 to 0.75; **fig. S6**), selected based on known target annotations and observed consistency in the compounds’ phenotypic effects in JUMP (**Methods, Fig. 3b**).

**Fig. 3:**
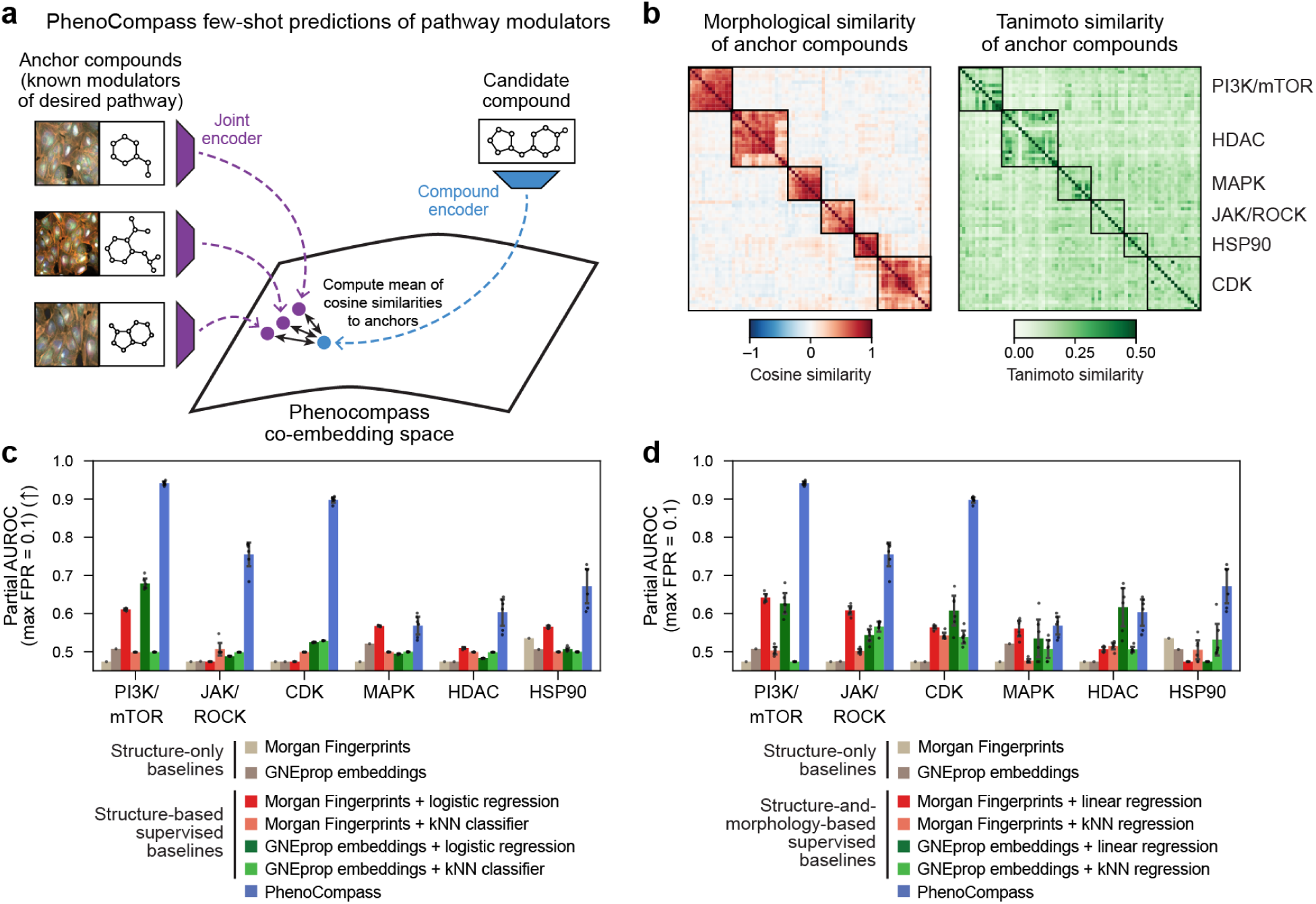
PhenoCompass few-shot predictions enable phenotypic virtual screening of structurally dissimilar pathway modulators. **a**, Schematic of the PhenoCompass framework for scoring pathway-specific chemical modulators using few-shot learning. For a particular pathway of interest, a set of known modulators serves as anchor compounds, which are embedded using the joint encoder. Pheno-similarity of candidate compounds is assessed as the mean cosine similarities of their compound embeddings to all anchor representations. **b**, Diversity of morphological similarities (left) and Tanimoto similarities (right) of the anchor compounds targeting the PI3K/mTOR, HDAC, MAPK, JAK/ROCK, HSP90, and CDK pathways. Heatmaps show pairwise cosine similarity and Tanimoto similarity (color bars) between anchor compounds. **c, d**, PhenoCompass accurately retrieves pathway modulators across diverse targets. Retrieval accuracy (y axis, pAUROC, max FPR = 0.1) of PhenoCompass, structure-only baselines, and baseline structure-based supervised classifiers (c) or baseline structure-and-morphology-based predictors (d) for identifying structurally dissimilar (Tanimoto similarity < 0.15) pathway modulators across six pathways (x axis). Bar colors indicate the specific representation and model combination used. Bars: mean ± 2 s.e. of six splits. Arrows: higher (↑) or lower (↓) values represent better performance.

We rigorously simulated virtual screens for each pathway using held-out JUMP data, ensuring the model had never encountered these annotated compounds during training (**Methods**). To explicitly test for generalization to novel chemical spaces (“scaffold hopping”), we evaluated the model’s accuracy in retrieving pathway modulators that were structurally dissimilar to the anchor set (TS < 0.15) in a leave-one-out fashion. We benchmarked PhenoCompass against two categories of baselines: (1) structure-only approaches, including structural similarity metrics and supervised classifiers trained on Morgan Fingerprints or pre-trained GNEprop embeddings to predict binary pathway labels; and (2) multimodal baselines explicitly trained to predict morphology features from compound structures, followed by a similarity lookup (**Methods**). For the PhenoCompass model, we used joint representations as anchors, as they tended to yield better retrieval performance compared to using only the compound or morphology representations (**fig. S7**).

PhenoCompass dramatically outperformed the baselines in identifying structurally novel modulators across most pathways, despite not being explicitly trained on pathway labels. For the PI3K/mTOR pathway, PhenoCompass reached a pAUROC of 0.94 (a 39% improvement over the next-best structure-based method), while CDK and JAK/ROCK retrievals achieved pAUROCs of 0.90 and 0.75 (70% and 49% improvements, respectively) (**Fig. 3c**). Notably, the majority of structure-based supervised baselines failed to perform better than random (pAUROC = 0.5), potentially due to the scarcity of positive training examples available for each pathway. PhenoCompass also decisively outperformed the multimodal predictive baselines (**Fig. 3d**), demonstrating relative improvements of 47% (PI3K/mTOR), 48% (CDK), and 24% (JAK/ROCK). One exception was the HDAC pathway, where a GNEprop linear regression baseline narrowly outperformed PhenoCompass. This was likely an artifact of the hydroxamate warhead shared across 14 of the 17 target compounds, which structure-based models could have exploited despite low overall structural similarity (**fig. S6d**). We also observed that the baseline models trained on both morphology features and compound structures consistently outperformed supervised baseline models trained on structure alone (**Fig. 3c, d**). Overall, these results demonstrate that multimodal representations, derived from abundant but unlabeled compound-phenotype pairings, can be successfully leveraged to discover structurally novel modulators of disease-critical signaling pathways.

### PhenoCompass accurately predicts binders of disease-associated proteins

To determine whether PhenoCompass could bridge the gap between phenotype-based and target-based drug discovery, we evaluated its ability to predict target-specific binding affinities despite being trained exclusively on broad morphological profiles. We scored millions of compounds from historical high-throughput screens (HTS) for PIK3CA, JAK3, and HDAC2 inhibition using the JUMP-trained PhenoCompass model as a few-shot learner on the relevant anchor sets from JUMP (**Methods**). The HTS data for each target consisted of: (1) a primary screen of up to two million compounds at a single concentration, and (2) a secondary screen, in which active compounds from the primary screen were re-tested across a range of doses to determine potency (pIC_50_) (**Fig. 4a**, **table S2**). Despite large distribution shifts between the JUMP and HTS chemical spaces (**fig. S8**), PhenoCompass’ pathway scores are robustly correlated with target potency (**Fig. 4b**). Active compounds (pIC_50_ ≥ 5) scored significantly higher than compounds that failed the primary screen (*p* < 10^−46^, one-sided Wilcoxon rank-sum test), and the model successfully distinguished the most potent inhibitors (pIC_50_ ≥ 7) from moderate ones (*p* < 10^−5^, one-sided Wilcoxon rank-sum test).

**Fig. 4:**
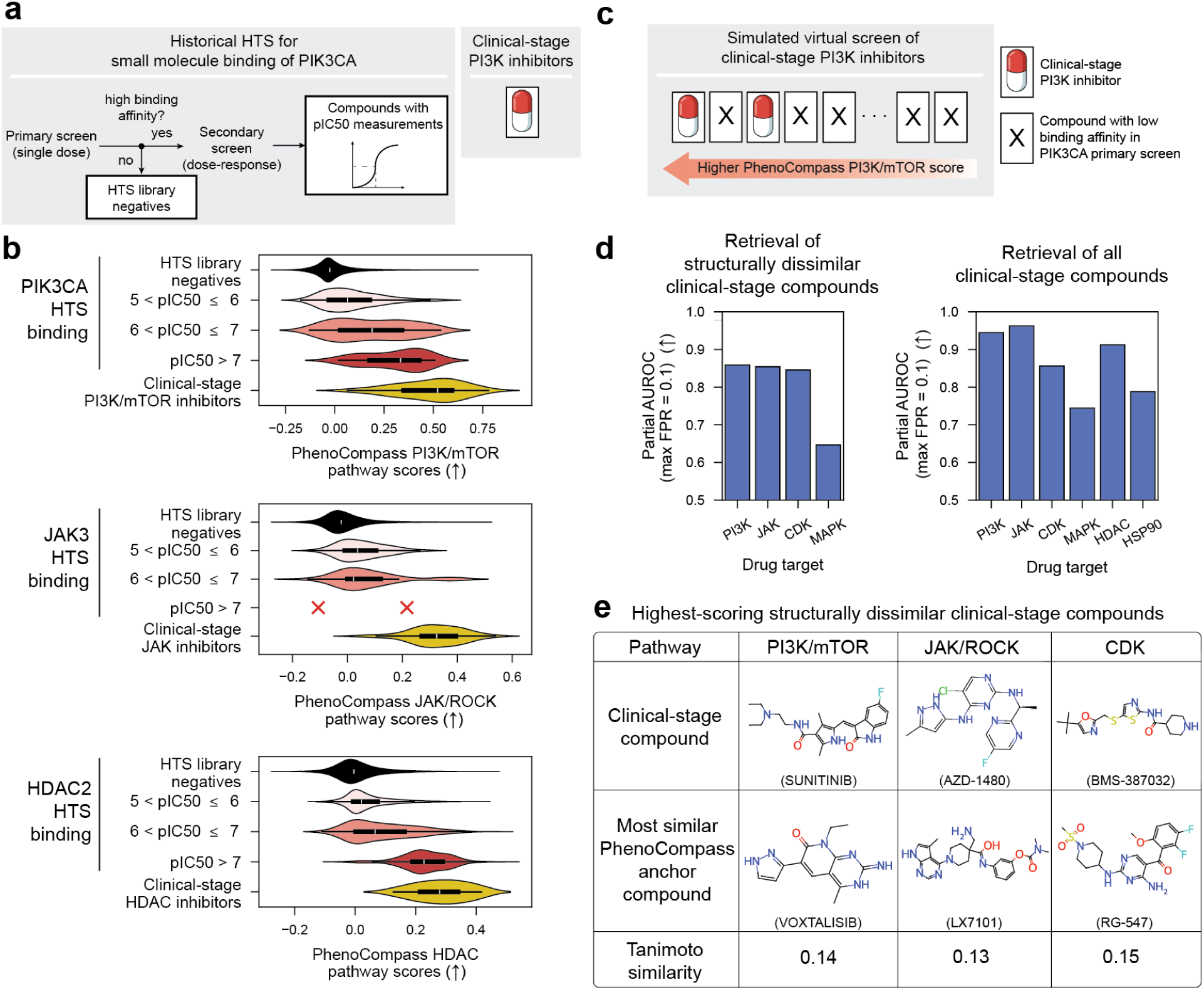
PhenoCompass predicts relative potency of HTS hits. **a**, Schematic of simulated virtual screen for PI3K inhibitors. Left: Historical high-throughput screening (HTS) data for PIK3CA binding consists of a primary screen of inhibition at a single dose, followed by a secondary dose-response screen for the active inhibitors to yield pIC_50_ measurements. Right: Known clinical-stage PI3K inhibitors constitute an additional ground truth hit category. **b**, PhenoCompass scores correlate with binding affinity. PhenoCompass pathway scores (x axis, violin plots) for different simulated HTS hit categories (y axis). PhenoCompass PI3K/mTOR, JAK/ROCK, and HDAC pathway scores were computed for the PIK3CA, JAK3, and HDAC2 HTS binding data, respectively (panels, top-to-bottom). **c**, Schematic of a simulated virtual screen of clinical-stage PI3K inhibitors using PhenoCompass. Clinical-stage PI3K inhibitors (ground truth hits) and compounds with low binding affinity in the PIK3CA HTS primary screen (ground truth negatives) are ranked by PhenoCompass PI3K/mTOR scores. **d**, PhenoCompass accurately retrieves clinical-stage inhibitors. Retrieval accuracy (y axis, pAUROC, max FPR = 0.1) of structurally dissimilar (Tanimoto similarity < 0.15) (left panel) and all clinical-stage compounds (right panel) in simulated virtual screens using PhenoCompass across different drug targets (x axis, bar plots). **e**, Chemical structures of the highest-scoring clinical-stage compounds (top row) that have low structural similarity (Tanimoto similarity < 0.15) to their corresponding most similar PhenoCompass anchor compounds (middle row) and their respective Tanimoto similarity (bottom row), across the PI3K/mTOR, JAK/ROCK, and CDK pathways (columns, left-to-right). Arrows: higher (↑) or lower (↓) values represent better performance.

In addition, we used PhenoCompass to score clinical-stage drugs not profiled in the JUMP dataset (**Fig. 4c**). These highly optimized therapeutics significantly outscored even the most potent HTS secondary screen hits (*p* < 10^−10^, all secondary screen compounds; *p* < 10^−6^ compounds with pIC_50_ ≥ 6; one-sided Wilcoxon rank-sum test). In a simulated virtual screen challenging the model to retrieve these clinical-stage drugs from a background of HTS compounds, PhenoCompass achieved remarkable predictive accuracy for PI3K-, JAK-, and CDK-targeting drugs, yielding partial AUROCs (max FPR = 0.1) of 0.86, 0.85, and 0.85, respectively, for compounds that are structurally dissimilar from the anchor compounds (TS < 0.15 to anchor sets) (**Fig. 4d**, **Methods**). Notably, the highest scoring set of structurally dissimilar PI3K, JAK, and CDK-targeting drugs have strikingly distinct structures compared to their most similar anchor compounds (**Fig. 4e**). These results point to PhenoCompass’s potential for discovering novel target binders without relying on any prior protein-ligand structural data.

### Prospective virtual screening identifies structurally novel, potent PI3K/mTOR inhibitors

We prospectively screened a 3.8-billion-compound subset of the Enamine REAL catalog^48^ for inhibitors of the PI3K/mTOR pathway^19–22^, using PhenoCompass to prioritize candidates at a scale far beyond conventional phenotypic screening, followed by systematic validation through assays of increasing phenotypic resolution^22^ (**Fig. 5a**). Bridging the large distribution shift between the JUMP and Enamine compound spaces presents a formidable computational challenge (**fig. S9**), yet it offers an unprecedented opportunity for novel chemotype discovery. After scoring the entire catalog for predicted PI3K/mTOR inhibition activity, we shortlisted the top 100,000 candidates, which outscored half of clinical-stage PI3K-targeting drugs (**fig. S10**), and applied stringent absorption, distribution, metabolism, and excretion (ADME) safety filters^49^ to purchase and synthesize a structurally diverse set of 396 “PhenoCompass compounds” (**Methods**).

**Fig. 5:**
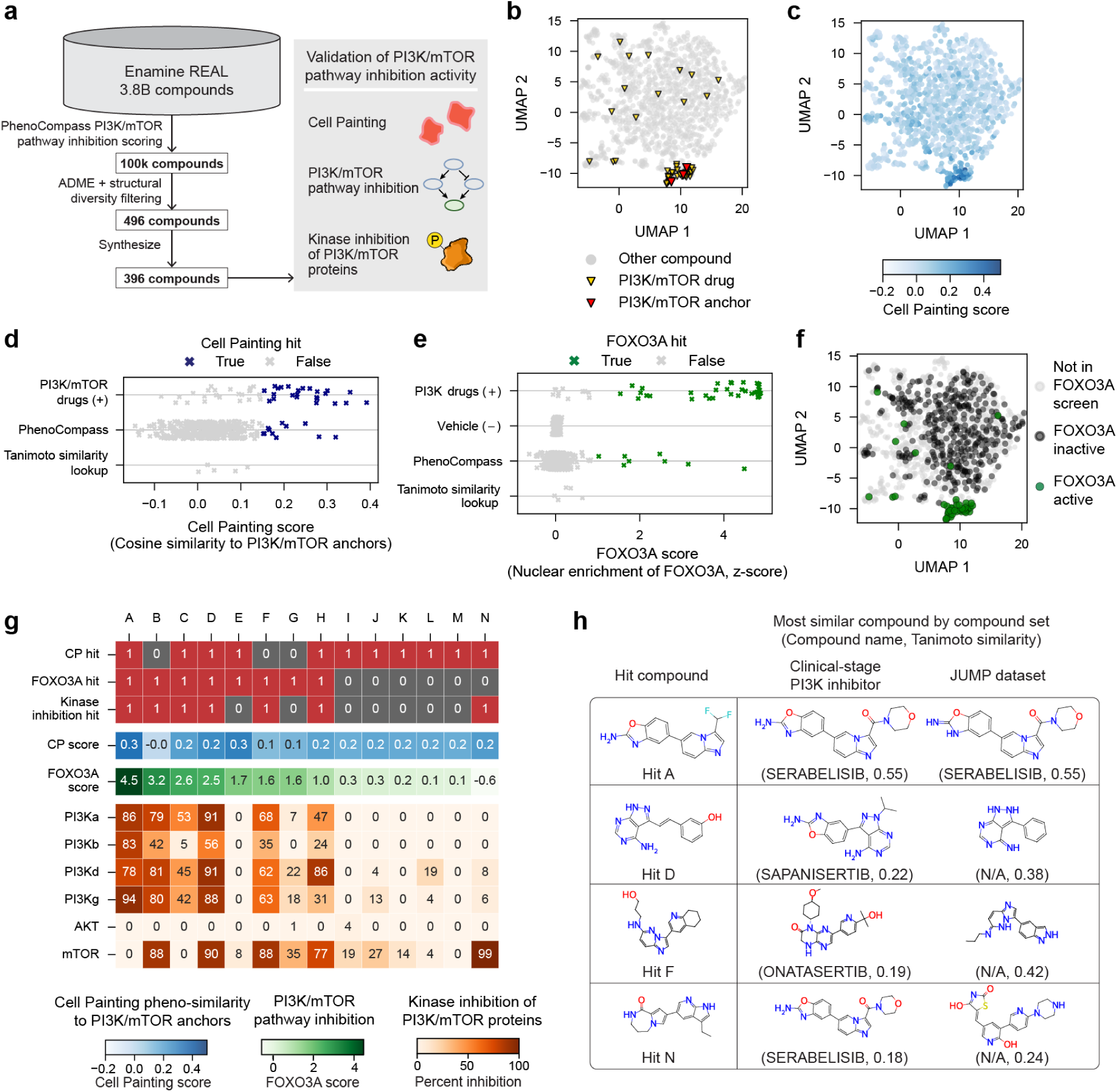
Large-scale virtual screening identifies structurally novel compounds with PI3K/mTOR inhibition activity. **a**, Workflow for performing large-scale phenotypic virtual screening for PI3K/mTOR pathway inhibitors using PhenoCompass. 3.8 billion Enamine REAL compounds are scored for PI3K/mTOR pathway inhibition, filtered by ADME properties and structural diversity, synthesized (left funnel), and subjected to three stages of experimental validation (Cell Painting, FOXO3A pathway signaling, and kinase inhibition; right panel) to yield confirmed hits. **b, c**, Cell Painting validation phenotypic map captures known PI3K/mTOR pathway modulators. UMAP visualizations of cell morphology profiles (markers) of compounds in the validation library (excluding plate controls), highlighting known PI3K- or mTOR-targeting drugs and PI3K/mTOR anchor compounds (b, color legend), as well as the resulting Cell Painting scores for each compound (c), computed as the mean cosine similarities of a compound to the PI3K/mTOR anchor compounds (color bar). **d**, Cell Painting scores (x axis) of PhenoCompass compounds and control compounds (y axis) relative to PI3K/mTOR anchor compounds. Cell Painting hits (blue crosses) are defined as compounds with Cell Painting scores greater than 0.15 (99th percentile of all control compounds) and *p* < 0. 01 relative to an empirical null distribution of Cell Painting scores derived from negative control samples (**Methods**). **e**, Nuclear enrichment of FOXO3A (x axis, z-score) for cells perturbed by PhenoCompass compounds and control compounds (y axis). FOXO3A hits (green crosses) are defined as compounds with z > 1 (**Methods**). **f**, FOXO3A hits cluster with known modulators in Cell Painting phenotypic space. UMAP visualization highlighting the FOXO3A hit compounds (green dots) against inactive (dark gray dots) and unscreened compounds (light gray dots). **g**, Summary of all Cell Painting, FOXO3A, and/or kinase inhibition-confirmed Enamine hits (columns A–N). Hits are ordered by their FOXO3A scores. Heatmaps show discrete hit status (top rows; red indicates a hit, gray indicates not a hit), quantitative Cell Painting and FOXO3A scores (middle rows; color bar; low, white; high, dark blue/green), and kinase inhibition percentages for specific targets (bottom rows; color bar; low, white; high, orange). **h**, Chemical structures of select hit PhenoCompass compounds alongside their most structurally similar clinical-stage PI3K inhibitor and their most similar compound in JUMP by Tanimoto similarity. N/A indicates that the compound has no common name.

As initial validation, we ran Cell Painting on U2OS cells to profile the morphological effects of the synthesized compounds together with a library of control compounds. This revealed a 33% bioactivity rate among the Phenocompass compounds (134 out of 396, **Methods**), compared to the 28% bioactivity rate in the JUMP dataset (30,082 out of 108,836), (**Methods**, **fig. S11a**). By projecting these profiles into a Cell Painting validation map (**Fig. 5b,c**), we found strong recapitulation of control compound MoAs (**fig. S11b**) and established a robust pheno-similarity metric for pathway inhibition that successfully recalled known PI3K/mTOR drugs (51% recall of mTOR/PI3K drugs and pAUROC (max FPR=0.1) = 0.77, **fig. S11c,d**). Applying a conservative hit-calling threshold, 11 of the 396 compounds (∼3%) exhibited specific PI3K/mTOR-like morphological signatures (**Methods**, **Fig. 5d**). This represents a 54-fold enrichment over the baseline hit rate (0.051%) of the full 108,836-compound JUMP dataset (**Methods**). Interestingly, many of the PhenoCompass compounds that did not register as PI3K/mTOR-related hits still showed significant pheno-similarity to other compounds (**fig. S11e**, **Methods**). In particular, the PhenoCompass compounds were enriched for inhibitors of PDGFR-beta and KIT, which are common off-targets of established PI3K and mTOR inhibitors^50^. In addition, we compared PhenoCompass against a baseline of 5 Enamine compounds with high structural similarity (TS > 0.5) to one or more of the PI3K/mTOR anchor compounds but were not among the highest scoring 100,000 PhenoCompass compounds. None of these baseline compounds exhibited bioactivity (“Tanimoto similarity lookup”, **Fig. 5d, fig. S11a)**.

As Cell Painting provides a broad readout of cell morphology, we orthogonally validated on-pathway inhibition using a FOXO3A nuclear translocation assay. FOXO3A is a downstream transcription factor that enriches in the nucleus upon PI3K/mTOR pathway blockade (**fig. S11f**)^51^. Screening our compounds under stringent enrichment thresholds yielded 8 FOXO3A hits, 5 of which directly overlapped with our Cell Painting hits (**Fig. 5e**, **Methods**). Crucially, the broad morphological signatures derived from Cell Painting were highly predictive of this targeted pathway readout: compounds inducing FOXO3A nuclear enrichment are co-clustered with PI3K/mTOR inhibitors in the Cell Painting validation map (**Fig. 5f**), and cosine-similarity to the PI3K/mTOR anchor compounds was a strong predictor of FOXO3A nuclear enrichment (pAUROC (max FPR=0.1) = 0.82, **fig. S11g**).

Finally, we profiled the 14 combined Cell Painting and FOXO3A hits (Hits A–N), together with five positive control compounds, against six key pathway kinases (PI3Ka, PI3Kb, PI3Kg, PI3Kd, AKT1, and mTOR) using targeted ADP-Glo biochemical assays (**Methods, fig. S11h**). Seven compounds were confirmed as direct kinase inhibitors (>50% inhibition) with diverse mechanisms of action (**Fig. 5g**). While several acted as pan-PI3K inhibitors, Hit N was highly mTOR-specific, exhibiting PI3K/mTOR inhibitor-like Cell Painting morphologies without triggering FOXO3A translocation. Most importantly, these confirmed inhibitors are also structurally diverse: only two shared a TS > 0.4 with any JUMP training compound, and internal similarity was exceptionally low (only one pair of hits had TS > 0.23) (**fig. S12a**). Compared to known agents, only two hits (A and H) exceeded TS > 0.3 to any PI3K/mTOR anchor, and only three (A, C, and H) exceeded TS > 0.3 to clinical-stage PI3K inhibitors (**fig. S12b**). While Hit A, the most potent phenotypic hit, structurally resembles the PIK3CA-specific inhibitor serabelisib^52^, Hits D, F, and N exhibited minimal similarity (TS ≤ 0.22) to any clinical counterpart (**Fig. 5h**). Furthermore, Hits D and F most closely align with sapanisertib and onatasertib, respectively, yet lack the strict mTOR selectivity of these investigational drugs^53,54^. Hit N proved the most structurally unique (TS = 0.18 to its most similar clinical-stage PI3K/mTOR inhibitor, serabelisib) and functionally distinct, demonstrating strong mTOR specificity rather than serabelisib’s PIK3CA selectivity. Overall, these diverse potent inhibitors provide highly valuable starting points for therapeutic development, showcasing the potential of virtual phenotypic screening at the pathway level with PhenoCompass.

## Discussion

Drug discovery campaigns are often forced to strike a difficult compromise between the ultra-high-throughput scale of target-based virtual screening and the physiologically relevant readouts of phenotypic screening. By developing PhenoCompass, we have introduced a framework that enables prospective virtual screening of ultra-large chemical libraries for complex, pathway-specific cellular phenotypes. We demonstrated PhenoCompass’s utility by training on over 100,000 compounds to identify novel inhibitors of the PI3K/mTOR pathway within the Enamine REAL library (3.8 billion compounds). We validated these hits across multiple biological resolutions spanning morphological profiles, pathway reporter activity, and direct kinase inhibition, achieving a 54-fold enrichment in hit rate relative to the training dataset. Our results demonstrate that self-supervised multimodal representation learning can unify high-dimensional Cell Painting data with small molecule structures, providing a powerful search engine for chemical space defined by biological states.

Prior approaches have been developed for virtual phenotypic screening; however, each has made different trade-offs between phenotypic resolution and chemical coverage. For instance, GNEprop^8^ successfully navigated large-scale libraries, but was limited to predicting low-dimensional phenotypes such as bacterial viability. Conversely, methods leveraging richer transcriptional readouts, such as DrugReflector^55^ and other models^56–60^ trained primarily on the LINCS^61^ dataset, offer high-dimensional profiles but have largely been restricted to drug repurposing within libraries of 10^5^ – 10^6^ compounds. Closely related to our work are methods that directly generalize from Cell Painting^45^, including multimodal contrastive learning frameworks like MoCoP^14^, CLOOME^15^, MIGA^16^, and MolPhenix^17^, yet these efforts have remained largely retrospective. A notable exception is PhenoModel^62^ which enabled a prospective screen over a comparatively small library of ∼335,000 compounds, validated by Cell Painting and viability across two cell lines. In contrast, PhenoCompass enables prospective, pathway-specific screening at billion-compound scale, supported by orthogonal phenotypic and biochemical validation.

Operationally, we view virtual phenotypic screening as complementary to target-based drug discovery. While PhenoCompass identified novel pathway modulators, these hits are unoptimized and require further medicinal chemistry optimization to achieve the potency, selectivity, and pharmacological properties required for clinical development^63^. Moreover, phenotypic screening identifies hits for a desired cellular state without explicitly identifying the underlying molecular target, relying on post-hoc deconvolution to elucidate precise molecular drivers and evaluate potential safety risks^64^. It is precisely this target-agnostic nature that allows high-dimensional PDD to excel at uncovering novel mechanisms that rational, single-target methods might overlook. As progress in experimental deconvolution techniques, such as mass spectrometry^65^ and genetic dependency screens^66^, continues to accelerate, the target identification bottleneck is likely to diminish, creating a critical opportunity for virtual phenotypic screening to rapidly prioritize biologically active chemotypes for downstream deconvolution. Furthermore, in oncology, efficacy often emerges from the simultaneous modulation of multiple nodes, causing diverse binding patterns to converge on a single morphological phenotype^67^. This is reflected in our results, where multiple activity profiles from structurally diverse candidates produced similar FOXO3A and Cell Painting signatures, ranging from pan-PI3K to mTOR-skewed inhibition, and compounds that did not meet our PI3K/mTOR criteria were enriched for commonly observed off-target kinases such as PDGFRβ and KIT.

Despite these advances, several limitations remain to be addressed. PhenoCompass was trained exclusively on the JUMP dataset, which relies on U2OS osteosarcoma cells. While many fundamental signaling cascades, such as the PI3K/mTOR pathway, are highly conserved across cell types, the discovery of modulators for tissue-specific diseases will require training on more complex biological model systems, such as primary cells, induced pluripotent stem cells, or organoids. Furthermore, PhenoCompass is trained only to predict morphological phenotypes through Cell Painting, whereas high-throughput transcriptomics provides a more readily integrated modality^68^, and may provide complementary signals for biological processes that are not detectable by imaging^69^. Transcriptomics could also provide a direct bridge to patient-derived state programs, for example by defining disease–healthy axes from scRNA-seq and prioritizing compounds that shift in vitro systems along the same trajectories^55^. Finally, PhenoCompass’s few-shot virtual screening framework heavily depends on the availability and phenotypic consistency of high-quality anchor compounds. For entirely novel biological pathways lacking known modulators, alternative querying strategies, such as using genetic perturbation anchors (e.g., CRISPR knockouts), will be necessary. Unfortunately, the multi-site origin and experimental design of the JUMP dataset introduced significant technical variation, preventing alignment between genetic and chemical spaces for this purpose.

Looking forward, as high-content profiling becomes more affordable, the data landscape for training virtual screening models will continue to evolve. Perturbation atlases will expand in both scale and diversity, evidenced by recent announcements of large consortia efforts, such as Illumina’s Billion Cell Atlas^70^, the Arc-Biohub-Tahoe Initiative^71^, and the Virtual Cell Pharmacology Initiative^72^. Increasing training set size by iterative, targeted data acquisition has led to major capability gains in other machine learning domains, including large language models^73^. Analogously, combining heterogeneous, large-scale consortia atlases and using lab-in-the-loop strategies^55^ to refine models could expand virtual phenotypic screening to a wider set of therapeutically relevant biological systems and phenotypic objectives. In this regard, frameworks such as PhenoCompass will serve as critical navigational tools, accelerating the exploration of high-content biology and the discovery of next-generation therapeutics.

## Materials and Methods

### Phenotypic Map Construction

#### JUMP Cell Painting Consortium Dataset

The JUMP Cell Painting dataset includes cell painting images from around 116,000 chemical and 22,000 gene perturbations on human U2OS osteosarcoma cells^18^. Data generation was split across 12 data-generating centers (or “sources”) with the following design: each compound has an average of 5 replicates from at least 3 sources. Cell painting data from each source are organized into batches and plates. In each plate, wells treated by DMSO served as negative controls. Between four and nine fields-of-view (FoVs) were imaged in each well. Each FoV consists of five channels: mitochondria (MitoTracker; Mito), nucleus (Hoechst; DNA), nucleoli and cytoplasmic RNA (SYTO 14; RNA), endoplasmic reticulum (concanavalin A; ER), and Golgi and plasma membrane (wheat germ agglutinin) combined with actin cytoskeleton (phalloidin) as ‘AGP’^18^. In addition to raw images, the dataset also includes morphological features extracted from a CellProfiler pipeline^18^. For our CellProfiler maps, we used these well-aggregated features.

### Image Representation Learning with DINO

As a preprocessing step before model training, illumination correction was applied to the Cell Painting images, and intensities of each channel were clipped at the 0.1st and 99.9th percentiles and scaled to the range [0, 1]. We then trained a DINO model^24^ with a cross-source sampling strategy^27–29^ to learn representations of cell morphological features. The DINO architecture includes a student branch and a teacher branch, and the model is trained by minimizing the cross-entropy loss between the embeddings produced by the two branches. With the cross-source sampling strategy, DINO takes image samples with the same compound treatment from different sources as input for student and teacher branches, respectively. In this way, the model is trained in a weakly supervised manner and is encouraged to learn morphological representations that are invariant to source-level batch effects. In line with Kim et al.,^25^ only FoVs from sources 2, 3, 6, and 8 in the JUMP dataset were used for training. Those FoVs have spatial dimensions ranging from 970 by 970 to 1024 by 1024. To take the advantage of the cross-source sampling technique, only perturbations with samples available in more than two sources were included.

During model training, random image crops of size 256 by 256 were sampled from the FoVs. Otsu thresholds were calculated in the DNA channel and only image crops with more than 1% foreground area in the DNA channel were selected for model training to avoid training on empty image crops. For each image crop, we further extracted one global crop (224 by 224) and four local crops (96 by 96) to enforce global-to-local consistency, which has been shown to improve the quality of DINO’s representation^24^. In addition, we applied a sequence of data augmentations including horizontal and vertical flips, brightness adjustments, and contrast adjustment to each crop. Finally, the images were standardized using the global mean and standard deviation over the intensities of all pixels across all images.

The ViT-small architecture was used with a patch size of 16 as the backbone, and a hidden dimension of 20,000 for the projector. We used a batch size of 128 and a learning rate of 0.002, with a 20-epoch linear warmup from zero followed by a cosine decay. The model was optimized using Adam for 200 epochs. We set the teacher temperature at 0.04, with a 20-epoch linear warmup from 0.01, and used a momentum of 0.996 for the teacher’s update. Gradient clipping was applied with a maximum gradient norm of 3.

During model inference, we divided each FoV into image tiles of 224 by 224 and fed these tiles into the model. The [CLS] tokens from the last four ViT blocks were concatenated to represent each tile. Tile embeddings were then averaged within each FoV to represent the corresponding FoV. Next, the representations of FoVs from the same well were averaged to build well-level embeddings.

### Feature post-processing and map construction

We applied similar post-processing pipelines to both DINO and CellProfiler features, with and without batch correction, to obtain four JUMP phenotypic maps. Unless mentioned otherwise, all post-processing steps apply to all four maps. Based on an evaluation of map quality when using different subsets of sources (see “Source filtering” below), we restricted the data to sources 1, 2, 3, 5, 6, 8, 10, 11. We dropped perturbations annotated as “Unknown”. We furthermore removed wells with fewer than ten cells in at least one image, or less than the expected image count per well (four images for sources 1 and 9; six images for sources 2 and 10; nine images for all other sources). Additionally, we dropped complete plates with fewer than 380 wells remaining post filtering (1470 remaining wells for sources 1 and 9). Following previous studies^14^, for the CellProfiler map only, we dropped CellProfiler features that are considered to be noisy or unreliable. We removed all inf and nan values in the data by discarding features containing more than 1% inf or nan and any remaining wells with inf or nan. We then normalized the data plate-wise using median and median absolute deviation, keeping only the features with variances between 0.1 and 5. Following this, CellProfiler features were reduced to 384 dimensions by PCA, while DINO features were kept at their full dimension of 1536. Finally, as we focused only on the compound perturbations, we generated canonical SMILES strings for each remaining compound perturbation using RDKit and removed compound perturbations for which canonicalization failed.

For batch-corrected maps only, batch correction was performed by typical variance normalization (TVN)^74^, and CellProfiler features for each well were additionally centered separately on the mean of all negative controls sharing the same “Metadata_platetype”.

### Phenotypic map evaluation

We assessed the quality of our phenotypic map using metrics for batch effect, replicate reproducibility and recall of compounds with shared target annotation. To estimate batch effect, we quantified the Kolmogorov-Smirnov (KS) statistic between the cosine similarities of 10,000 randomly sampled well-level profile pairs from the same batch and a size matched distribution randomly sampled across all batches. Similarly, we calculated replicate reproducibility as the KS statistic of 10,000 randomly sampled well-level profile pairs with the same perturbation ID and a size matched random sample. To evaluate the recall of compounds with shared target annotation, we leveraged target gene and MoA annotations from OpenTargets as ground truth. We largely followed the evaluation approach outlined in Celik et al.,^30^ and quantified recall of compound pairs with a shared target gene or MoA annotation within the 10% most extreme pairs of perturbation-level profiles (5% highest and 5% lowest cosine similarity).

### Identifying bioactive perturbations from Cell Painting images

To assess the rate of bioactive perturbations, we compute a perturbation phenotype significance compared to random background on the batch-corrected DINO map. We closely follow the perturbation consistency metric described by Celik et al.^30^ Specifically, for each compound, we collected all well-level morphological profiles for the compound, computed the mean cosine similarity of each well-level profile to all of the remaining well-level profiles for the compound, and computed the mean of these values, which we refer to as the leave-one-out (LOO) cosine similarity. An empirical null distribution of LOO cosine similarities was generated by randomly sampling across all compounds the same number of well-level profiles as available for the compound of interest, computing a LOO cosine similarity for this sample of profiles, and repeating this procedure 9,999 times. The mean LOO cosine similarity for a compound was compared against this sample size-matched empirical null distribution to assess the statistical significance of a compound’s activity. We converted p-values to false discovery rates (FDR) using Benjamini-Hochberg correction, and compounds with *FDR* < 0. 01 were considered bioactive. Using this procedure, we found bioactivity rates of 28% (30,082 of 108,836) and 33% (134 of 396) in the JUMP and Cell Painting validation datasets, respectively.

For genetic perturbations, we calculated bioactivity rates based on a separate map build that included both compound and genetic perturbations. Post-processing steps were the same as in “Feature post-processing and map construction” with the following changes: We used plate-wise sphering instead of TVN and centered by negative control wells in each perturbation modality separately. We report this metric separately for expressed and unexpressed genes. Unexpressed genes are defined as genes with less than -3 zFPKM in expression data of HeLa cells obtained from DepMap^75^ (**table S1**). Stratifying bioactivity in this way revealed very high rates among all genetic perturbations, including CRISPR KO perturbations of unexpressed genes (95%). This coincides with a large fraction of perturbations that always co-occur in the same well position across technical replicates/plates (94.9%), prompting us to dismiss genetic perturbations for inclusion in our final map due to concerns of a significant amount of their morphological heterogeneity being driven purely by technical confounders rather than true underlying biological signal.

### Source filtering

We hypothesized that restricting our map to a subset of all available sources could improve map quality due to the heterogeneity between data sources^23,25^. To identify the optimal subset of sources, we analyzed batch effects, replicate reproducibility and compound target prediction performance for single sources and all source pairs. For this analysis, we processed the full dataset (all sources, including genetic perturbations) using the pipeline described in “Feature post-processing and map construction”, except for the use of negative control sphering instead of TVN and centered by negative control wells in each perturbation modality separately (same map build as in “Identifying bioactive perturbations from Cell Painting images”). We further subset this map to positive control plates (TARGET2) with fixed plate layout and perturbation composition across all sources, ensuring a fair comparison, and we restrict to single sources and source pairs for evaluation. These plates were recorded by all sources except source 1, which was excluded from this analysis. Batch effect and replicate reproducibility were quantified as described above. To quantify compound target prediction performance, we leveraged compound target annotations included in the TARGET2 plate metadata and calculated the mAP of retrieving perturbations with the same target annotation. This analysis (**fig. S1a-c**) indicates that sources 4, 7, 9 and 13 are outliers that are harder to integrate with the other sources (sources 7, 13), display relatively poor replicate reproducibility (source 9) or target prediction mAP (sources 4, 9).

We furthermore quantified recall of OpenTargets MoA annotations (as described above) for multiple source selections (**fig. S1d**): all sources, sources using confocal microscopes (sources 2, 5, 6, 8, 10, excl. source 7, 13), sources used by Kim et al.,^25^ (sources 2, 3, 6, 8) as well as all sources excluding outlier sources (sources 1, 2, 3, 5, 6, 8, 10, 11). In the latter case, we included source 10 despite its relatively poor performance on the target prediction task, as well as source 1 which didn’t record TARGET2 plates. Both sources are members of the “Wave 1” group of sources that exchanged nominated perturbations with each other^18^, and their inclusion improves the overall replicate structure in the data. Prior to recall calculation, each of these data subsets was processed separately as described (except for using negative control sphering instead of TVN) and the final recall metric was averaged over 2, 4, 6, 8, 10% most extreme pairwise perturbation-level profiles. Based on these results, we used sources 1, 2, 3, 5, 6, 8, 10, 11 for the final DINO map and PhenoCompass training.

### Map visualization

UMAP visualizations of the final JUMP phenotypic map were constructed using the umap-learn library with the following parameters: n_neighbors=25, min_dist=0.2, negative_sample_rate=40, repulsion_strength=1.5, metric=”cosine”. For the UMAP visualization in **Fig. 1e**, we show the morphology profiles of (1) compounds associated with the top 10 MoAs in the JUMP phenotypic map and (2) compounds that were determined to be bioactive (see “Identifying bioactive compounds for Cell Painting datasets” above) and that had strong effects (embedding L1 norm ≥ 6).

### Curating JUMP compound annotations and pathway-anchor selection

In order to identify tight clusters of compounds with overlapping annotations in the JUMP phenotypic map, we first curated compound annotations across multiple resolutions and ground-truth datasets. We considered the following three resolutions: target gene (e.g. MEK1, MEK2), target gene family (e.g. MEK), and target gene functional class (e.g. Serine-Threonine Kinase). First, we combined annotations from an in-house database (N=833 compounds) and OpenTargets (N=964 compounds; N=384 unique MoAs), mapping target genes using data from the NIH National Library of Medicine. Next, we added data from ZINC15 and ChEMBL to annotate additional compounds, using an affinity cutoff of 100nM to assign target genes. The full matrix of compounds and their annotations (N=1239) are provided in **Data S2.**

To identify clusters of compounds with known MoAs, we applied the following procedure: First, we selected a subset of phenotypically diverse compounds from the set of all annotated compounds. We then computed all pairwise cosine similarities between the DINO representations of this subset. Next, we constructed a graph, connecting two compounds with an edge only if the cosine similarity between their DINO representations was in the top 0.1% of all pairwise cosine similarities. For each connected component in the graph, we excluded outliers that are connected to fewer than 25% of the other compounds in the component. We also retained only the connected components with at least 5 remaining compounds for use as seed clusters. Then we expanded these seed compound clusters by connecting them to annotated compounds if their cosine similarities were within the top 0.5% of all pairwise cosine similarities computed above. Finally, we re-applied the same outlier as before, dropping outlier compounds that are connected to fewer than 25% of other compounds within these expanded clusters, resulting in a final set of 136 annotated compounds across 14 components. Mechanisms of action for these final compounds are then assigned the gene-target level of annotation in order of priority: in-house, OpenTargets, ChEMBL, and ZINC15.

From these compounds, we selected a smaller subset to use as anchors for few-shot predictions of pathway-specific effects. We focused on the PI3K/mTOR, JAK/ROCK, CDK, HDAC, MAPK, and HSP90 pathways, as compounds targeting these pathways and/or target classes demonstrated distinct clustering in the JUMP phenotypic map. We further refined the compound clusters associated with these pathways to arrive at our final sets of anchor compounds for each pathway. We retained subclusters of compounds with high phenotypic similarity and consistent gene target annotations, excluding subclusters containing promiscuous compounds.

### PhenoCompass Computational Framework and Retrospective Validation

#### Molecular features

We used multiple molecular featurization strategies to encode compounds into a fixed embedding space for downstream processing and to assess similarity, including molecular fingerprints and deep-learning-derived features. For fingerprints, we computed Morgan Fingerprints (2048 bits, radius = 3), Avalon fingerprints (1024 bits), Extended-Reduced Graph (ErG) fingerprints, and a set of 200 molecular features^7^ using the 2023.9.5 version of RDKit. These features were generated using RDKit’s default parameters and any reference to these fingerprints assumes these parameters unless otherwise stated. Tanimoto similarities were computed between Morgan fingerprints of tautomer-canonicalized structures, using RDKit’s TautomerEnumerator class and TanimotoSimilarity function, respectively. For deep-learning-derived embeddings, we used a GNEprop model that was pre-trained in an unsupervised manner as in the original publication^8^, using the same parameters as for the full PhenoCompass architecture (GNEprop (pre-trained), **table S3**).

#### PhenoCompass architecture

To enable phenotypic virtual screening of chemical compounds, we developed the PhenoCompass framework. It leverages geometric multi-modal contrastive (GMC) representation learning to align modality-specific representations and joint representations comprising information across all modalities^39^. Specifically, GMC consists of modality-specific encoder networks, a joint modality encoder network, a projection network for each of the modality-specific and joint modality encoders, and a common encoder network (**fig. S3**).

We use two distinct models to embed compounds. The first, “PhenoCompass (GNEprop)”, uses a graph-neural-network-based compound encoder that takes as input a molecular graph and employs the GNEprop^8^ architecture, which leverages Graph Isomorphism Networks and Jumping Knowledge Networks^76,77^. The second, “PhenoCompass (Curated Features)”, uses a multilayer perceptron (MLP) compound encoder and operates on a vectorized collection of the following curated molecular features and fingerprints:, GNEprop (pre-trained), Morgan Fingerprints, Avalon fingerprints, ErG fingerprints, and a set of 200 molecular features computed with RDKit (see “Molecular features” above). We use batch-corrected and compound-level aggregated DINO features as input as morphology embeddings.

In the following, we outline the PhenoCompass neural network architecture. We denote individual paired morphology and compound datapoints with subscript *i* and different parts of the architecture with the superscript letters *E* (embedding), *P* (projection), *C* (compound), *M* (morphology), and *J* (joint). First, for a compound *i*, the compound encoders generate an initial compound embedding 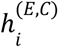 and the morphology encoder (implemented as an MLP) maps aggregated DINO embeddings to an initial morphology embedding 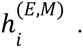 The joint encoder *i* (*E*,*M*) uses the same modality-specific encoder architectures, but with independent weights. These encoders produce embeddings 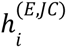 and 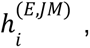 respectively, that are concatenated to yield an initial joint embedding 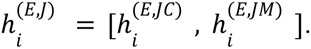 Each of the initial compound, morphology, and joint embedding then serve as input to their respective projection networks, implemented as MLPs that yield compound, morphology, and joint projection embeddings 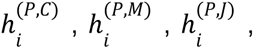 respectively. Finally, a single common encoder network (MLP), takes each of these projection embedding as input to generate final representations for each of the compound, morphology, and joint modalities 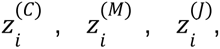 respectively, which we refer to as compound representations, morphology representations, and joint representations.

#### Contrastive loss

Each of the PhenoCompass models were trained using the GMC objective function^39^, which focuses on geometrically aligning a model’s joint representations with each of their matching compound and morphology representations. GMC frames geometric alignment as a classification task for pairs of representations, in which positive pairs are encouraged to be similar and negative pairs are encouraged to be dissimilar. Because GMC aims to align joint representations with each of their corresponding compound or morphology representations, each positive pair of representations consists of one joint representation and its matching compound or morphology representation. All other pairings within and between the joint and alternate modalities are defined to be negative pairs.

Specifically, we consider a mini-batch *B* with representations 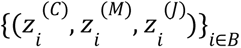 and write 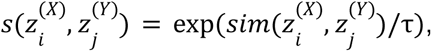 where *sim*(*u*, *v*) represents the cosine similarity between vectors *u* and *v*, τ is a temperature hyperparameter, and *X*, *Y* ∈ {*C*, *M*, *J*} denote modalities as above. For a sample *i*, we define 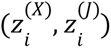 and 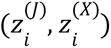 to be the positive pairs for all *X* ∈ {*C*, *M*}. The corresponding contrastive loss associated with these positive pairs is the cross-entropy loss,

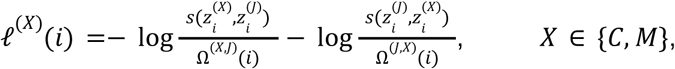

Where Ω^(*X,Y*)^(*i*) denotes a normalization factor over the positive pair and the background of negative pairs,

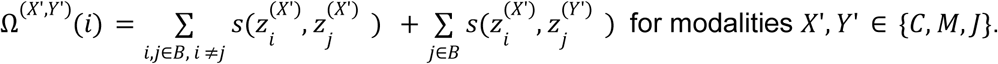

Finally, these per-sample loss terms are combined across the samples of each mini-batch,

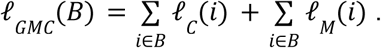

#### Consistency regularization loss

For the GNEprop PhenoCompass model, we further adapted consistency regularization ^78^ to the multimodal setting. Compared to regular contrastive learning, where similarity of a positive pair is enforced relative to a background of negatives, consistency regularization further enforces that the distributions of similarities to the negatives align between the representations of a positive pair. In other words, for an example positive pair 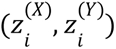 and a negative representation 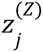 for 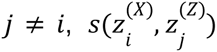 and 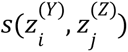 are encouraged to be similar. We implement this as follows: For a datapoint *i* and a pair of modalities *X*, *Y* ∈ {*C*, *M*, *J*}, we define the similarity distribution to the negatives as the vector 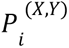 with entries

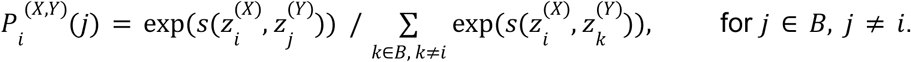

The alignment between the distributions for different query modalities is then quantified with the symmetrized Kullback-Leibler divergence (also known as Jeffreys divergence),

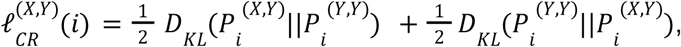

where 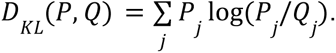. Summing over the full batch, we obtain

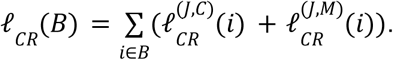

When training the GNEprop PhenoCompass model, we combine both GMC loss and consistency regularization loss with a hyperparameter α

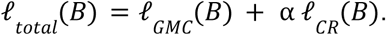

#### Cross-modal retrieval and few-shot virtual screening with PhenoCompass

To perform cross-modal retrieval with PhenoCompass, candidate embeddings for morphology 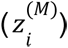 and compound structure 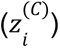 are obtained and then compared using cosine similarity in the same representation space, with top-rated pairs serving as predicted matches (**Fig. 2a**).

To infer compounds for pathway-specific effects, PhenoCompass uses few-shot learning: For a specific pathway, a set of known modulators of this pathway are first selected as anchors. These anchors are embedded with PhenoCompass, using either its morphology, compound, or joint encoder to generate these embeddings. To query a candidate compound for the desired pathway-specific effect, we first embed it using PhenoCompass’s compound encoder. We then score this candidate compound by computing the mean of the cosine similarities between its representation and the representations of each of the anchors (**Fig. 3a**).

#### Data splitting

To assess the ability of PhenoCompass to generalize to novel classes of compounds, we used scaffold-cluster splitting^8^ to generate training, validation, and test splits for the JUMP dataset. Scaffold-cluster splitting involves encoding scaffolds of compounds using a pre-trained GNEprop model, connecting representations for these scaffolds via a kNN graph (*k* = 5) using Euclidean distance as a distance metric, and then identifying clusters on this graph using Leiden clustering^79^ with resolution *r* = 200. Each of the resulting scaffold-clusters and all of their constituent compounds were treated as a distinct unit for splitting the JUMP dataset, such that compounds with structurally similar scaffolds would not appear in the same training, validation, or test sets.

We split the JUMP dataset such that the set of curated annotated compounds (see *Compound annotations*) and compounds within the same scaffold-clusters as this set were apportioned to the test set. We then sampled entire scaffold-clusters uniformly from the set of remaining scaffold-clusters until the number of compounds associated with these sampled scaffold-clusters reached or exceeded 5% of the number of compounds in the JUMP dataset. These compounds were assigned to the validation set. All remaining compounds were assigned to the training set. We generated six scaffold-cluster splits using this sampling procedure, with the random seeds used for sampling scaffold-clusters matching those used for initializing weights for each model. The test set consisted of 6,945 compounds. Across different scaffold-cluster train-test splits, the validation and training sets ranged from 5,452 – 5,497 compounds and 96,394 – 96,439 compounds, respectively.

These initial splits were used to train and evaluate models for cross-modal retrieval (**Fig. 2**) and few-shot prediction (**Fig. 3**) within the JUMP dataset. For experiments with HTS data (**Fig. 4**) and the large-scale PI3K/mTOR virtual screen (**Fig. 5**), we added the test sets into the training set while retaining the same validation sets. We then re-trained models on these new splits with the same hyperparameters as in the initial splits to maximize efficient use of the available data.

The UMAP visualization of scaffold-clusters in the JUMP dataset (**fig. S4a**) was generated by constructing a kNN graph (*k* = 5) of the GNEprop scaffold representations using Euclidean distance as a distance metric and then applying the scanpy implementation of UMAP using default parameters.

#### Model training and ensemble learning

Models were trained using the hyperparameter and training configurations indicated in **table S3**. Model training was stopped when the plate-matched Hits@5 metric computed on the validation set failed to increase over 10 epochs. The model with the highest Hits@5 metric was retained (see “Evaluating cross-modal retrieval”).

For pathway activity predictions (**Fig. 3**–**5**), we further leveraged ensemble learning. Specifically, we trained an ensemble of 12 PhenoCompass models, 6 using the PhenoCompass (GNEprop) model and the other 6 using the PhenoCompass (Curated Features) model. The model architecture and hyperparameters within each class of models were kept the same, while the weights for each model were initialized with different random seeds. For the simulated pathway-level virtual screens within JUMP (**Fig. 3**), we performed few-shot predictions using the mean of the few-shot prediction scores across only the two PhenoCompass (GNEprop) and PhenoCompass (Curated Features) models for each random seed in order to assess seed-to-seed variability in the results. For the final PhenoCompass ensemble of models (**Fig. 4**, **5**) we used the mean of the few-shot prediction scores across all 12 models as the pathway activity score.

#### Evaluating cross-modal retrieval

To evaluate a model’s ability to accurately retrieve a compound given a related image of cells perturbed with that compound (and vice versa), we compute Hits@K values. These values indicate the proportion of compound-image pairs for which, given an image (compound), the model can retrieve the matching compound (image) within the top K highest scoring compounds. We uniformly sampled 99 mismatched compounds (images) for each image (compound) to serve as the background. Evaluations on the validation set involved exclusively sampling background compounds from the validation set, and evaluations on the test set sampled only from the test set. The results reported in **Fig. 2** were calculated on the test set.

This regular Hits@K metric could potentially be inflated due to confounding, where a compound-image pair is solely recovered due to shared technical effects. To address this, we designed a modified Hits@K metric to account for plate-level batch effects. We refer to this metric as the *plate-matched Hits@K*. Specifically, for each well-level image for a compound, we one-hot encode the plate IDs corresponding to these images and add these one-hot encoded representations across images to generate a vector of plate ID counts per compound. The similarity of the plate ID vectors between two compounds indicates the similarity of the distribution of plates from which their image data was collected, with higher degrees of similarity serving as a proxy for higher shared batch effects. We used the L1 distance to assess similarity between two plate ID vectors. For the image-compound (compound-image) retrieval task, instead of randomly sampling background compounds as above, for each image (compound), we used the 99 mismatched compounds (images) with the highest plate ID similarity to that of the image (compound). Samples for which fewer than 99 mismatched compounds have L1 distance less than 0.5 were discarded from these evaluations, as they represent samples for which we cannot sufficiently generate plate ID-matched backgrounds.

We compared PhenoCompass’s performance with linear regression and *k*-nearest neighbors (kNN) regression (*k* = 10) models. These models were trained on either Morgan Fingerprints (see “Molecular features”) or pre-trained GNEprop embeddings of compounds to predict a compound’s DINO representations directly using least squares loss. Nearest neighbors for the kNN models were determined using Tanimoto similarity and Euclidean distances for the Morgan Fingerprints and GNEprop (pre-trained) embeddings, respectively.

#### Predicting bioactive compounds in the JUMP dataset

To evaluate the utility of different molecular representations for predicting bioactive compounds (**Fig. 2d**), we train supervised classifiers for compound bioactivity using the same set of train-test scaffold-cluster splits as above. Binary labels for bioactivity were obtained as all compounds with *FDR* < 0. 01 following the test described in “Identifying bioactive compounds from Cell Painting images” above. We fit either Logistic Regression or kNN classifiers (*k* = 10) to embeddings from PhenoCompass’s compound encoder, Morgan Fingerprints, or GNEprop (pre-trained) embeddings. Nearest neighbors for the *k*NN models were determined using Euclidean distance for the pre-trained GNEprop embeddings models and cosine similarity for the Morgan Fingerprints and PhenoCompass compound embeddings models. The logistic regression model was trained with “balanced” class weights and default elastic net regularization hyperparameters in scikit-learn.

#### Simulating virtual screens within JUMP

To assess the retrieval of compounds targeting a specific pathway (**Fig. 3c,d**), we first identified the anchor compounds we curated for the pathway (see “Curating JUMP compound annotations and pathway-anchor selection” above). We simulated the retrieval of each compound in this anchor set by leaving out a single compound and employing the remaining compounds in the set as few-shot examples. The retrieval task entailed prioritizing this left-out compound against a background of compounds that consisted of all compounds in JUMP for which we did not have annotations and that had dissimilar phenotypic effects from all of the anchor compounds.

For the structure-only baseline methods, we scored each candidate compound as its mean structural similarity to the anchor compounds, using Tanimoto similarity and Euclidean distance for the Morgan Fingerprints and GNEprop embeddings, respectively. For the supervised structure-based baseline methods, anchor compounds were labeled as positives and all remaining compounds in the training set from the scaffold-cluster splits as negatives, respectively. The logistic regression model was trained with “balanced” class weights and default elastic net regularization hyperparameters in scikit-learn. Nearest neighbors for the *k*NN (*k*=100) models were determined using Euclidean distance for GNEprop (pre-trained) embedding models and cosine similarity for the Morgan Fingerprints and PhenoCompass compound embeddings models. For the structure-and-morphology-based baseline models, we used the same baseline models as for cross-modal retrieval (see “Evaluating cross-modal retrieval” above). Few-shot predictions for a compound were made using these models by first predicting the morphology profiles of all compounds. The predicted morphology profile of a compound was then scored by computing the mean cosine similarity of its predicted profile with the predicted profiles of the anchor compounds.

For each set of anchor compounds, we evaluated retrieval accuracy by computing the mean pAUROC with a max FPR of 0.1 across the compounds within the set. To assess performance in the more challenging case of structurally dissimilar compounds (**Fig. 3d**), we restricted the query anchor compounds for each ground truth compound to those with a Tanimoto similarity of less than 0.15 to the ground truth compound, dropping compounds for which this set was empty.

#### Simulating virtual screens of HTS hits and clinical-stage compounds

To simulate virtual screens beyond the JUMP dataset, we compiled data from historic in-house high-throughput screening (HTS) assays of the Roche/Genentech HTS library for specific targets as additional test sets (**Fig. 4a,b**). These screens were performed in two stages: an initial primary single-point inhibition screen at 5µM, and a secondary hit-confirmation assay in dose-response format to generate IC_50_ values for selected primary-screen positives. The assays for each target were as follows. PI3K: recombinant PI3K-alpha/p85 activity was assayed using a fluorescence polarization assay to detect conversion of phosphatidylinositol(4,5)-phosphate to phosphatidylinositol(3,4,5)-phosphate (Echelon Biosciences). HDAC2: compounds were screened using a TR-FRET immunodetection assay of K382-acetylated p53-derived peptide (CisBio). JAK3: compounds screened using the ADP-glo method to detect phosphorylation of poly-(Glu4Tyr1) peptide (Promega Biosciences). Compounds were categorized based on their outcomes in the primary and confirmation screens: “HTS library negatives” if they were not selected for the confirmation screen, and pIC_50_ bands (5 < pIC_50_ ≤ 6, 6 < pIC_50_ ≤ 7, 7 < pIC_50_) according to their confirmation screen results if they were selected.

To leverage PhenoCompass in this setting, we used the full 12-model ensemble whose training data included the annotated anchor compounds. (see “Data splitting” and “Model training and ensemble learning”). Based on their annotations, we selected the PI3K/mTOR, JAK/ROCK, and HDAC pathway anchor sets to score the compounds in the PIK3CA, JAK3, and HDAC2 assays with PhenoCompass, respectively. We used the joint representations for all anchors in these sets, taking advantage of both the compound structure information and their corresponding Cell Painting images in JUMP.

To curate compounds with likely even stronger potency and efficacy than HTS hits, we extracted compounds from ChEBML with documented affinities to our anchor annotations that had been investigated clinically, meaning they had likely undergone additional medicinal chemistry optimization (e.g. “Clinical-stage PI3K inhibitors”, **Fig. 4a**). Specifically, using the ChEMBL web-client, we first extracted all Target ChEMBL IDs associated with a keyword search of the anchor annotation (PI3K: “PI3-Kinase”, JAK/ROCK: “Janus kinase” and “Rho-associated protein kinase”, HDAC: “Histone deacetylase”, HSP90: “HSP90”, MAPK: “Mitogen-activated”, CDK: “Cyclin-dependent kinase”). Target ChEMBL IDs associated with the keyword were filtered to cases containing a “Target Type” of either “Single Protein”, “Protein Family”, or “Protein Complex” with >10 documented IC50-based bioactivity profiles for the organism “Homo Sapiens”. For each anchor annotation, compound bioactivities from each of the associated Target ChEMBL IDs were concatenated and only the named compounds (clinical or post-clinical stage) were accepted. Finally, a manual curation step removed named-compounds where the bioactivity association was thought to be too weak or low confidence.

To assess virtual screening performance, we selected the set of clinical-stage compounds related to each anchor set as positives. For PI3K, JAK, and HDAC, negatives were defined as compounds from the related primary screen that failed to progress to the secondary screen. For CDK, MAPK, and HSP90, which lacked related primary screens, we used the negatives from the HDAC2 screen, as it was the largest available HTS dataset. (**Fig. 4c–e**). Compounds already present in the JUMP dataset were excluded when evaluating retrieval accuracy, as these compounds were used during training of the PhenoCompass models. Before computing Tanimoto similarities between JUMP and ChEMBL compounds, we apply tautomer canonicalization using RDKit’s TautomerEnumerator class to both sets to ensure consistent structural comparison. For each of the PI3K/mTOR, JAK/ROCK, and CDK pathway anchor sets, we visualized the highest scoring clinical-stage compound that shows specificity for and is clinically documented as a modulator for target(s) in the corresponding pathway (**Fig. 4e**).

### Prospective Discovery and Validation

#### Virtual Screening of PI3K/mTOR inhibitors on the Enamine REAL Database

##### PhenoCompass Predictions

We first downloaded a snapshot of the Enamine REAL database and filtered it by applying the following cutoffs on basic molecular descriptors: molecular weight between 250 and 450 Da, topological polar surface area (TPSA) between 60 and 120, calculated LogP (cLogP) between 1 and 4, number of H-bond acceptors between 0 and 8, number of H-bond donors between 0 and 4, and number of rotational bonds between 0 and 8, obtaining roughly 3.8B compounds and their SMILES strings. We then used PhenoCompass to score each of these compounds for PI3K/mTOR pathway inhibition activity. We retained the 100,000 highest-scoring compounds, as the 100,000th-ranked compound achieved a PhenoCompass score that exceeds the scores of roughly half of known PI3K-targeting drugs (**fig. S10**).

##### Compound ADME and structural diversity filtering

We used the multi-task neural network model from Napoli et al. (2025)^49^ to predict ADME (absorption, distribution, metabolism, and excretion) properties for all of the PhenoCompass-predicted compounds. Compounds predicted to exhibit undesirable ADME properties were removed from consideration to enrich for compounds that would pass safety criteria in drug development. Specifically, we excluded compounds with functional groups known to be reactive or that pose safety/selectivity risks, compounds with low predicted cell permeability, and compounds with high predicted cellular efflux^80^. To do so, we used a sphere exclusion clustering procedure, in which we iteratively select the compound with the highest PhenoCompass score and exclude from consideration any remaining compounds within a Tanimoto similarity radius of 0.45 from the selected compound. In total, we ordered 495 compounds from Enamine, of which 396 were effectively synthesized.

#### Cell Painting validation

##### Cell Culture and Harvesting

U2OS cells were sourced from Genentech’s Cell Bank (ATCC HTB-96). Cells were maintained in RPMI-1640 (Gibco) supplemented with 10% (v/v) fetal bovine serum (Gibco) at 37 °C, 5% (v/v) CO2, 95% humidity. All cells were pre-banked at passage 5 before screening. Cells were collected for experiments at about 70% confluency in culture flasks. Cells were first dissociated from culture flasks using the TrypLEExpress enzyme (Gibco) and resuspended in full growth RPMI media. Cells were counted using a Vi-CELL (Beckman Coulter), and then diluted in full growth RPMI to achieve a count of 50000 cells/ml (2500 cells/well when dispensed at 50 μL per well). The cell suspension was dispensed into PhenoPlate-384 Ultra microplates (Revvity) using a Multidrop Combi with standard-tube cassette (ThermoFisher). After cells were dispensed, microplates were left at room temperature on the stable lab bench surface for 1 hour to ascertain uniform cell attachment in the well, before incubation at 37 °C, 5% (v/v) CO2, 95% humidity. All plates were incubated for 24 hours before compound perturbation.

##### Compound Perturbation

In addition to the PhenoCompass compounds, we assayed the following control compounds: 47 PI3K/mTOR-targeting drugs, 57 plate control compounds (selected for phenotypic effect diversity), and a reference set of 736 target-annotated literature compounds. All compounds were stored at 10 mM stock concentrations in DMSO. In the compound perturbation step, compounds were plated in 384-well Echo Low Dead Volume (384 LDV) qualified source plates (Beckman Coulter: 001-16128). Then, after cells were attached to the assay plate for 24 hours, compounds were delivered to the assay plates using Echo 555 acoustic dispenser (Labcyte). For Cell Painting assay, all compounds were assayed at 10 uM final concentration, with 5 replicates per perturbation in assay plates. All assay wells were back-filled with DMSO to make sure that all assay wells contain 0.1% (v/v) DMSO final concentration. All perturbation assignments were randomized across plates to minimize plate position and batch effect as much as possible. After compound addition, assay plates were incubated at 37 °C, 5% (v/v) CO2, 95% humidity, for 24 hours.

##### Cell Staining

After cells were incubated with compounds for 24 hours, Cell Painting staining procedure was performed according to 2023 updated protocol by Cimini et al.^81^ with minimal adjustment as follows:

1. Replacing Hanks’ balanced salt solution (HBSS) washing solution with Phosphate Buffered Saline (PBS)
2. Cell Painting staining solution was filtered sterile using a 0.22 um filter (Corning, CLS430517) before adding to fixed cells, to minimize plate contamination after long-term storage

##### Imaging

Cells were imaged with Opera Phenix (Revvity, USA) using a 20× air objective lens, widefield, with 2×2 binning setting. 9 fields of images were collected per well, which results in about 2500 cells/well.

##### Feature extraction

Images were exported from Opera Phenix as 16-bit tif files (1280 x 1280 pixels). Images were analyzed using CellProfiler (version 3.1.9) using the same pipeline as in the JUMP paper^18^. The nuclei segmentation was performed by CellPose^82^ based on the DNA channel. Subsequent cytoplasmic segmentation was done in CellProfiler, based on the RNA channel. All cells touching the boundary of the image were excluded from analysis. A total of about 4700 features were calculated for each cell. Features can be roughly categorized as (1) intensity; (2) colocalization measurements; (3) granularity; (4) textural measurements of objects; (5) neighboring objects; (6) distribution of staining intensity patterns; (7) size/shape metrics.

##### Constructing the Cell Painting validation phenotypic map

We constructed a series of phenotypic maps from the computed CellProfiler features via a grid-search of different feature normalization, dimensionality reduction, and linear batch correction procedures (**table S4**). In particular, feature normalization was performed per plate, using all of the wells associated with the negative control compounds and the set of plate control compounds that were included in each plate. Batch effect reduction, reproducibility, and biological recall were quantified in a similar manner to described above in “Cell Painting Image Representation Learning”, except that in-house annotations for target genes were used to assess biological recall. We selected the phenotypic map that ranked most highly for a combination of batch effect reduction, reproducibility, and biological recall metrics. UMAP visualizations of the Cell Painting validation map were constructed using the umap-learn library with the following parameters: n_neighbors=25, min_dist=0.1, negative_sample_rate=40, repulsion_strength=1.5, metric=“cosine”.

##### Calling PI3K/mTOR Cell Painting hits

We defined pheno-similarity to the PI3K/mTOR anchor compounds as the mean cosine similarity of a compound’s morphological profile to those of the anchor compounds. We restricted the anchor compounds to three out of a total of four anchor compounds present in both the JUMP dataset and the Cell Painting validation data, excluding one anchor compound that had markedly lower phenotypic similarity to the remaining anchor compounds in the validation dataset compared to JUMP (**fig. S11c**). Hits were defined as those compounds whose PI3K/mTOR similarity exceeded the 99th percentile of pheno-similarity scores for all non-mTOR/PI3K-targeting control compounds present in both JUMP the validation data, and that achieved *p* < 0. 01 in the following statistical test: Each compound’s pheno-similarity score was compared with a plate-matched empirical null distribution of 10,000 pheno-similarity scores for the negative controls. In turn, each of the scores in the null distribution was computed by sampling a negative control well from each plate containing the compound being assessed, taking the mean of the negative control wells’ morphological profiles, and then computing pheno-similarity with respect to this mean profile.

To compute comparable hit rates in the JUMP dataset, pheno-similarity to the PI3K/mTOR anchor compounds was defined analogously, with the only difference that the empirical null distributions were constructed using samples from all compounds in a plate instead of only the negative controls. This was done since more substantial batch effects caused the morphological profiles for the negative controls (DMSO) in this dataset to not be consistently centered at the origin. For consistency with the Cell Painting validation data, we computed pheno-similarity with respect to the same three PI3K/mTOR anchor compounds described above and excluded all 13 PI3K/mTOR anchor compounds from the hit rate calculation, as these anchor compounds were curated based on their similarity to each other.

##### Evaluating phenotypic effects of non-hit PhenoCompass compounds

For each PhenoCompass compound that did not show PI3K/mTOR inhibition activity, we identified the most morphologically similar annotated control compound. For this set of most morphologically similar control compounds, we then assessed the enrichment of the control compounds’ target genes relative to the frequency of occurrence of the target genes across all control compounds. Statistical significance was assessed via a hypergeometric tests for each drug target, and drug targets with *p* < 0. 05 were considered enriched.

#### FOXO3A translocation assay

Recombinant U2OS cells stably expressing human FKHRL1 (FOXO3) (GenBank Acc. NM_001455) fused to the N-terminus of enhanced green fluorescent protein (EGFP) were purchased from Thermo Fisher (cat# R04-009-02).

##### Cell culture and treatment

Cell culture and compound addition were as described above. Compounds were tested as a 12-point dose response curve (starting concentration 10 uM, 3-fold dilution). After 1 hour incubation, cells were fixed by formaldehyde to a final concentration of 6.25% v/v for 30 min at room temperature before washing 3x with PBS, then 20 ul of nuclear staining solution (5µg/ml Hoechst 33342 in PBS) was added. Plates were then incubated for 30 minutes at room temperature, washed 3x with PBS, sealed and imaged.

##### Imaging

Cells were imaged with Opera Phenix (Revvity, USA) using a 20× air objective lens, widefield, with 2×2 binning setting using 405nm ex/435-480 em and 488 nm ex/500-550 nm em channels. 9 fields of images were collected per well, which results in about 2500 cells/well.

##### Feature extraction

Images were exported and segmented as described earlier. For FOXO3A translocation assay, only GFP intensity in nuclei, cytoplasm and cell were extracted.

##### Calling FOXO3A reporter assay hits

We computed the per-cell enrichment of FOXO3A nuclear translocation as the base-10 logarithm of the ratio of the mean fluorescence intensity in the nucleus to the cytoplasmic ring region. Per-cell ratio values were robustly z-scored using the median and median absolute deviation (MAD) of all cells in the negative control wells from the same plate. They were then clipped to be within -5 and 5, and successively aggregated to per-well and per-compound z-scores by arithmetic mean. Statistical significance of a compound’s FOXO3A nuclear translocation was computed by comparing the per-compound z-score with a plate-matched empirical null distribution of 1 million negative control scores, computed by sampling a negative control well from each plate containing the compound being assessed and then taking the arithmetic mean of the negative control wells’ z-scores. We converted p-values to false discovery rates (FDR) using Benjamini-Hochberg correction. A compound was considered a hit if its *FDR* < 10^−4^ and per-compound z-score > 1, resulting in calling 35 of 47 (74%) of positive control PI3K-targeting drugs as hits.

#### Kinase inhibition assays

Compounds were tested for inhibition of selected lipid and protein kinases by Reaction Biology Inc. (Malvern PA). PI3K kinases were tested using the ADP-Glo^TM^ assay and AKT was assayed by the HotSpot™ method under standard conditions as described by the manufacturer. Compounds were tested at 10 µM and 1 µM, in the presence of 10 µM ATP. Inhibition is expressed as the mean of two replicates normalized to percent of uninhibited (DMSO) control. Capivasertib, AZD-8835, Dactolisib, Apitolisib, and Idelalisib were used as positive control compounds (**fig. S11f**). A compound was considered a kinase inhibition “hit” if it inhibited the activity of at least one of the target kinases by at least 50%. To compare kinase inhibition hits with JUMP and clinical-stage compounds, we apply tautomer canonicalization to all three sets of compounds using RDKit’s TautomerEnumerator class before computing Tanimoto similarities.

## Acknowledgments

The authors gratefully acknowledge Colin Grambow for facilitating access to the Enamine dataset, and Henri Dwyer and Joseph Kleinhenz for support with compute.

## Author contributions

T.B., D.R., and J.-C.H. conceptualized the study. P.H., H.Y., B.H., and O.M. performed data curation and processing for the JUMP dataset. H.Y., B.H., J.-C.H., and D.R. developed the DINO methodology; H.Y. and B.H. implemented the software. A.W., H.Y., B.H., and G.G. performed formal analysis and visualization for this stage. A.W., H.Y., Z.L., G.S., D.R., J.-C.H., and T.B. developed the PhenoCompass methodology; A.W., H.Y., and Z.L implemented the software. G.G., J.M., and A.W. performed data curation for compound annotations and HTS resources. A.W., H.Y., G.G., and Z.L. led formal analysis and visualization. N.S., Z.L, G.S., and A.W. performed Enamine library curation. N.S. developed the methodology and software for compound filtering. N.K. conducted the wet-lab investigation, supervised by J.M. A.W., N.K., J.-C.H., D.R., and J.M. developed the validation methodology; N.K., A.W., and J.M. curated the data. A.W. led the formal analysis and visualization. G.S., J.M., J.-C.H., D.R., and T.B. provided supervision and resources. A.W., J.-C.H. and D.R. wrote the original draft of the manuscript; all authors reviewed and edited the final text.

## Data, code, and materials availability

Code, pre-trained models, phenotypic maps, and validation data from the Cell Painting experiments, FOXO3A nuclear translocation assays, and kinase inhibition assays conducted at Genentech related to this study are publicly available at https://github.com/Genentech/PhenoCompass and archived on Zenodo (https://doi.org/10.5281/zenodo.20367744). The following pre-trained models are included: PhenoCompass co-embedding ensemble (12 model checkpoints with 6 random seeds each for combined fingerprint and consistency regularization architectures); GNEprop structure encoder; and precomputed MoA anchor embeddings for PI3K/mTOR, HSP90, JAK/ROCK, HDAC, CDK, and MAPK pathways. We used the following publicly available datasets: Cell Painting morphological profiles from the JUMP-CP consortium (cpg0016-jump) available on the Cell Painting Gallery (https://registry.opendata.aws/cellpainting-gallery/)^18^; compound annotations from ChEMBL 33 (https://www.ebi.ac.uk/chembl/)^83^, Open Targets Platform (https://platform.opentargets.org/)^84^, and ZINC (https://zinc.docking.org/)^85^.

---

**Fig. S1:**
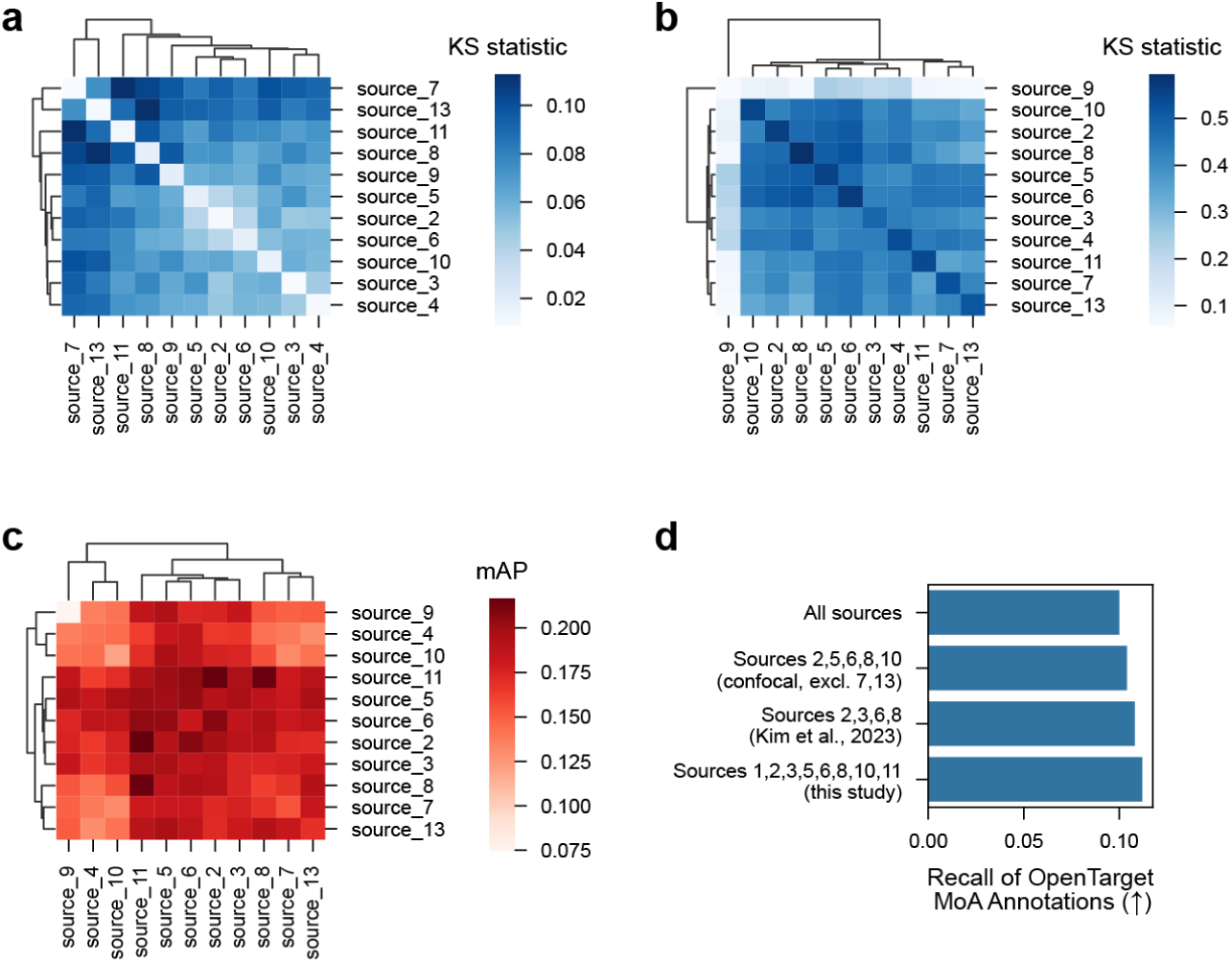
Data source quality varies with respect to batch effects, reproducibility, and target prediction. **a–c**, Pairwise source (off-diagonal) and single source (diagonal) metrics for batch integration based on CellProfiler features (**a**, KS statistic on batch variable; color bar: low, blue; high, red), replicate reproducibility (**b**, KS statistic on perturbation variable; color bar: low, blue; high, red), and compound target prediction (**c**, mean average precision (mAP); color bar: low, blue; high, red). Metrics are computed using TARGET2 plates to ensure fixed plate layout and perturbation composition across all sources (**Methods**). Source 1 did not record this plate type and was excluded from the analysis. Four sources exhibit either high batch effects (sources 7, 13), low replicate reproducibility (source 9), or low target prediction performance (sources 4, 9). **d**, Recall of OpenTargets MoA annotations (x axis) for specified data subsets (y axis). Included subsets are all twelve sources, all confocal sources excluding 7 and 13 (due to high batch effect), the four sources used by Kim et al. (2023), and the eight sources used in this study. Values reflect the mean over 2, 4, 6, 8, and 10% similarity cutoffs. Arrows: higher (↑) or lower (↓) values represent better performance.

**Fig. S2:**
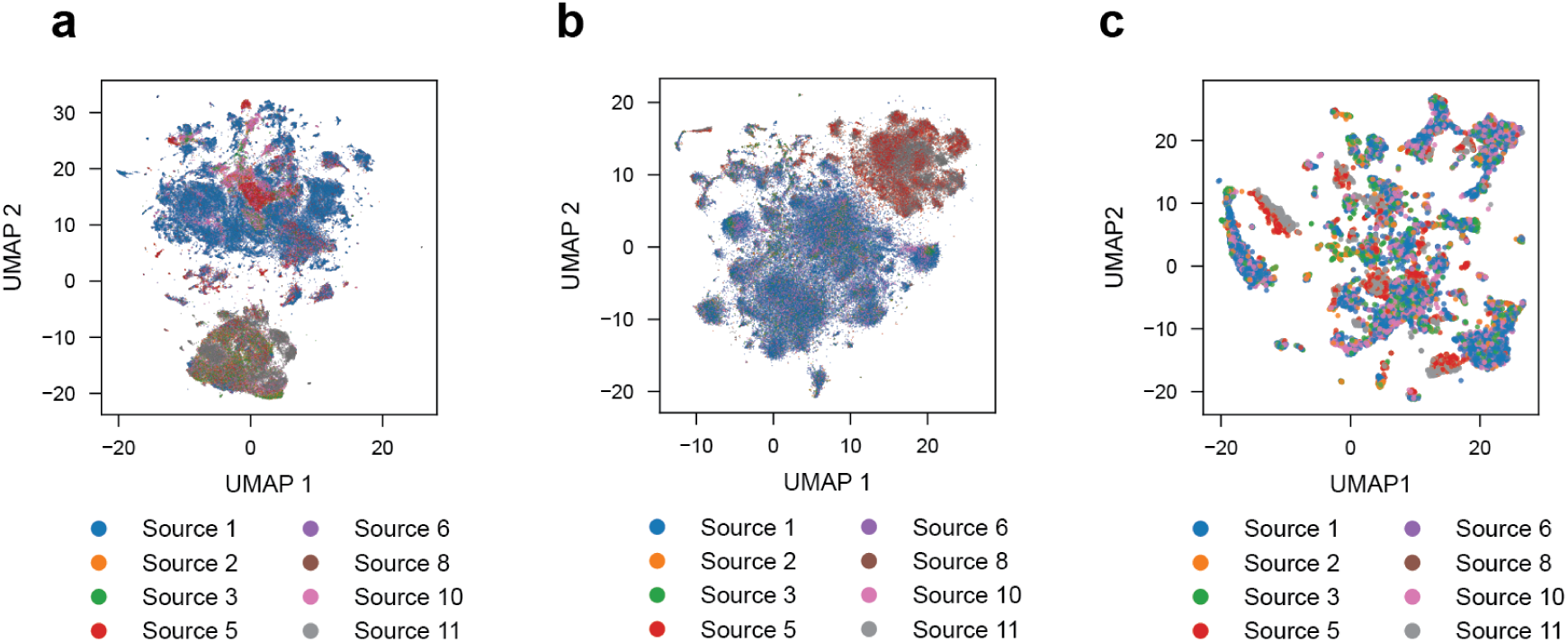
UMAP visualizations of source-related batch effects. **a–c,** Uniform Manifold Approximation and Projection (UMAP, scatter plots) of mean compound-level morphological profiles (dots) using the DINO phenotypic map without batch correction (a), with batch correction (b), with batch correction and subsetted to compounds with strong effect, as in Fig. 1e (c; **Methods**). Compounds are colored by their most frequently represented source (color legend).

**Fig. S3:**
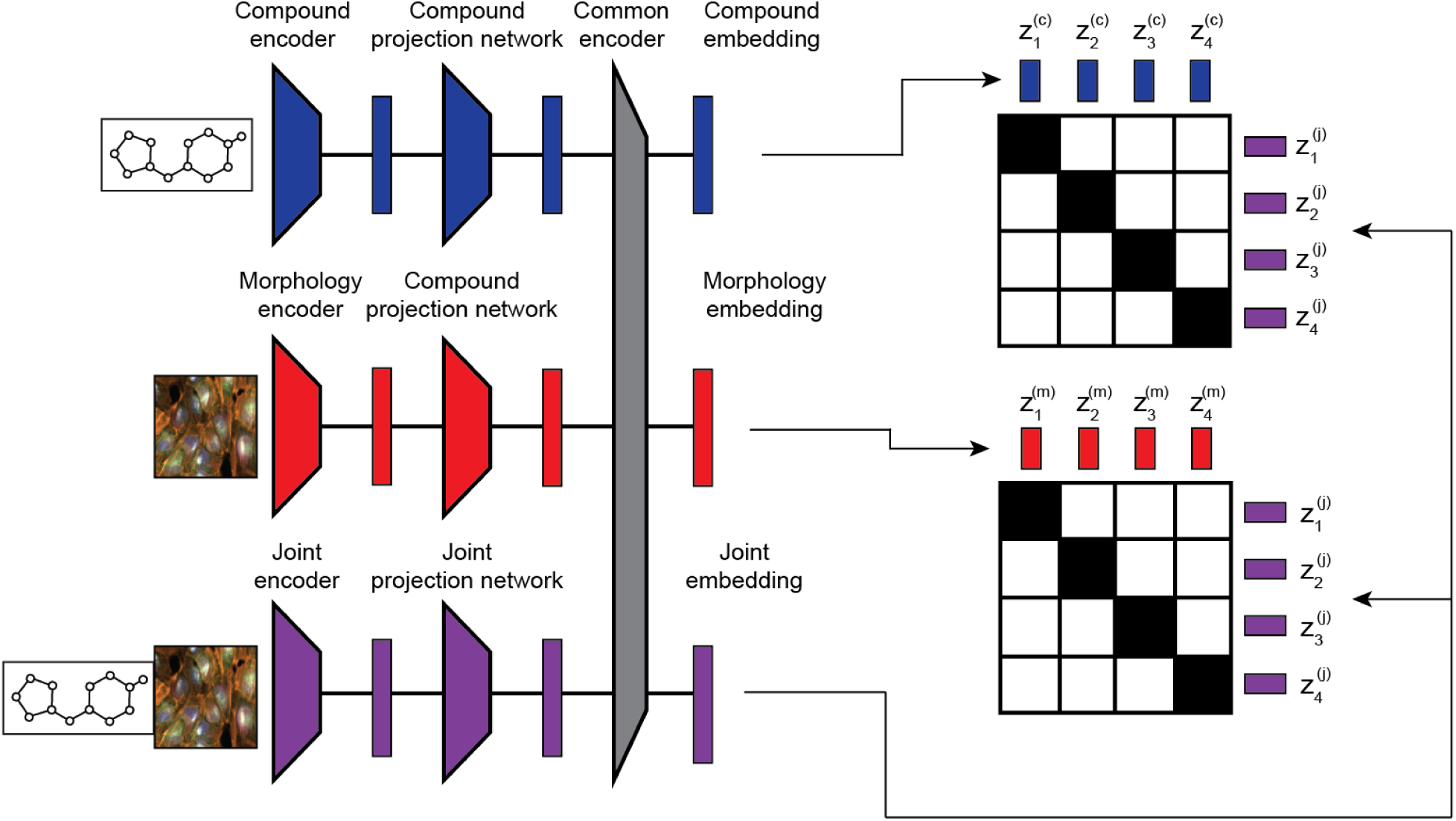
Schematic of the PhenoCompass geometric multimodal contrastive (GMC) learning architecture. PhenoCompass learns compound, morphology, and joint representations. Modality-specific inputs (left) are processed by compound and morphology encoders (center) and their respective projection networks to create modality embeddings. Simultaneously, a joint encoder produces embeddings from both inputs. Geometric alignment is achieved by enforcing contrastive loss between the compound/morphology representations and the joint representations (right).

**Fig. S4:**
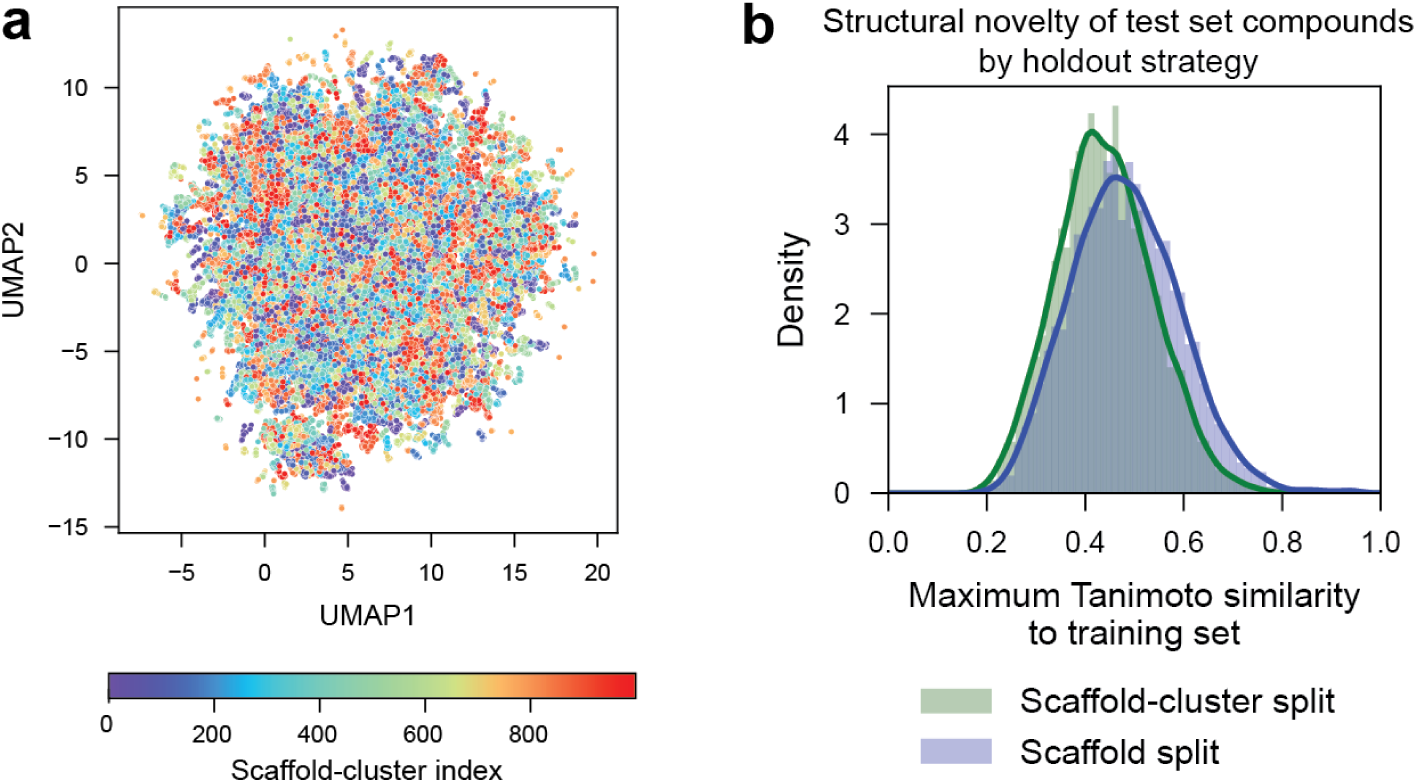
Training set properties. **a**, UMAP visualization (scatter plot) of pre-trained GNEprop embeddings for the molecular scaffolds of compounds in the JUMP dataset. Colors represent distinct groups of compounds in the same scaffold-clusters. **b**, Density distributions (y axis) of the maximum Tanimoto similarity of test compounds to the training set (x axis) when using scaffold-cluster split (green) versus conventional scaffold split (blue).

**Fig. S5:**
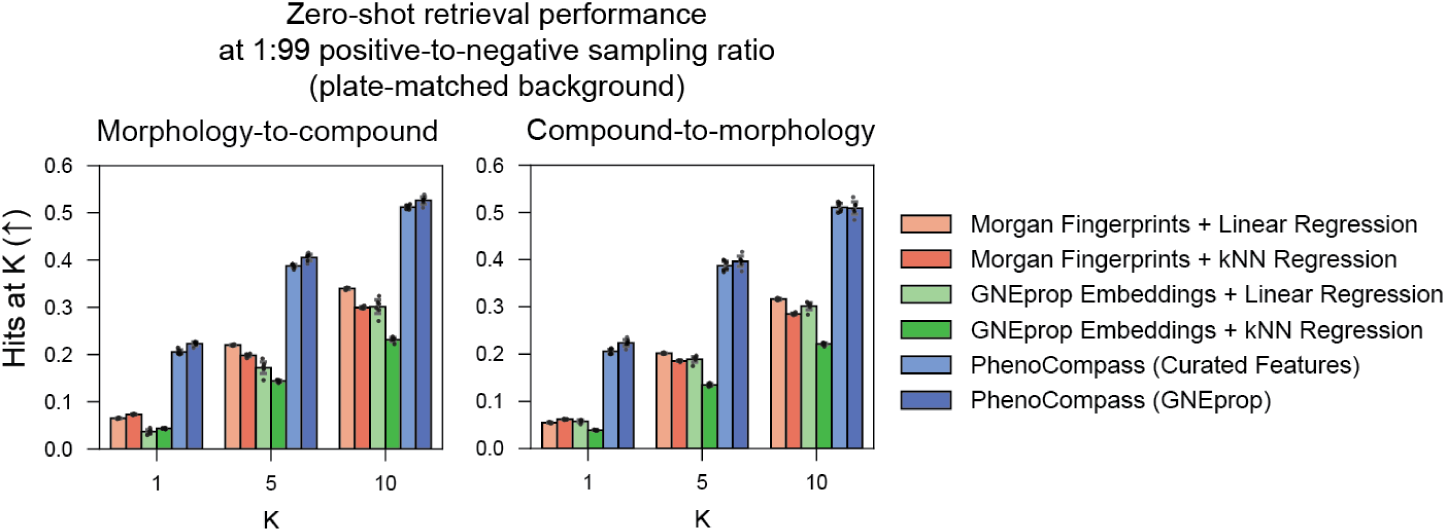
PhenoCompass zero cross-modal retrieval performance is robust to plate confounding. Accuracy of PhenoCompass and baseline models for the morphology-to-compound (left) and compound-to-morphology (right) retrieval tasks using plate-matched backgrounds to account for potential plate confounding (**Methods**). Retrieval performance (y axis) is evaluated as Hits at K for K = 1, 5, and 10 (x axis) across varying models (bar colors).

**Fig. S6:**
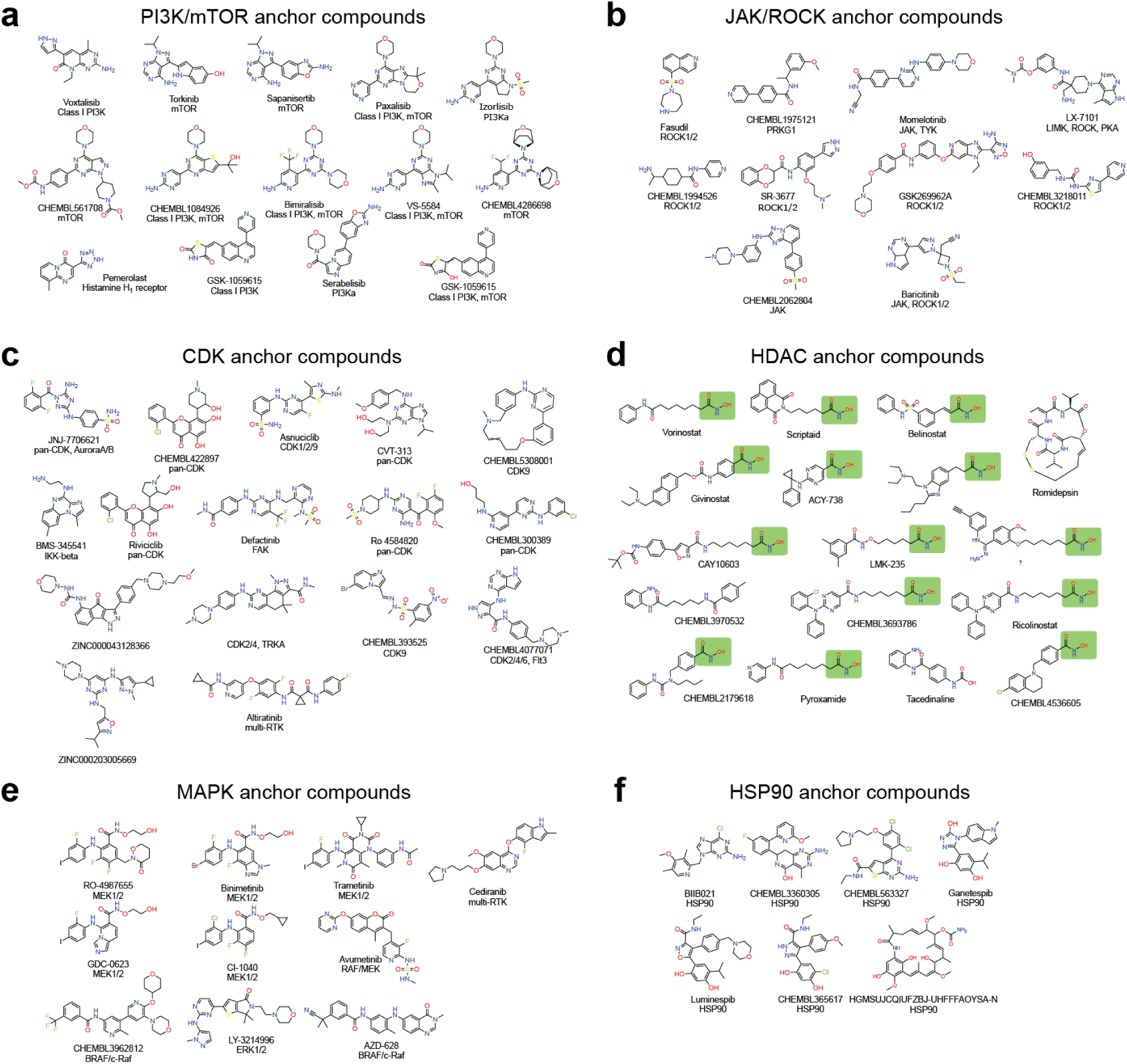
Molecular structures of anchor compounds. **a–f**, Chemical structures and associated target annotations for compounds in the PI3K/mTOR (a), JAK/ROCK (b), CDK (c), HDAC (d), MAPK (e), and HSP90 (f) pathway anchor sets. The green boxes in (d) indicate a hydroxamate warhead that is common to the majority of the compounds in the HDAC anchor set.

**Fig. S7:**
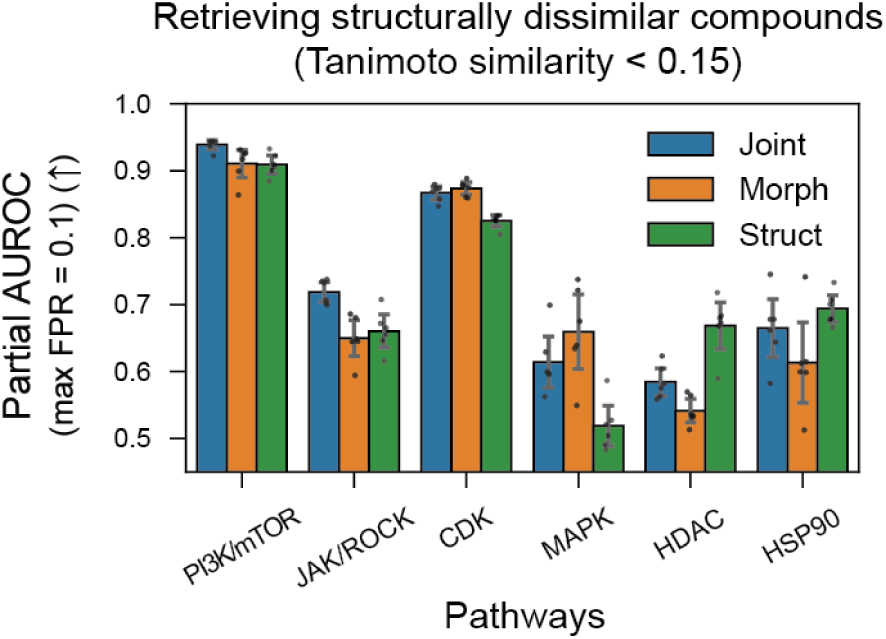
Comparison of PhenoCompass retrieval accuracies using different anchor representation types. Retrieval accuracy (y axis, partial AUROC, max FPR = 0.1) for identifying structurally dissimilar (Tanimoto similarity < 0.15) pathway modulators across various pathways (x axis) using joint, morphology, or structural representations as anchors (bar colors).

**Fig. S8:**
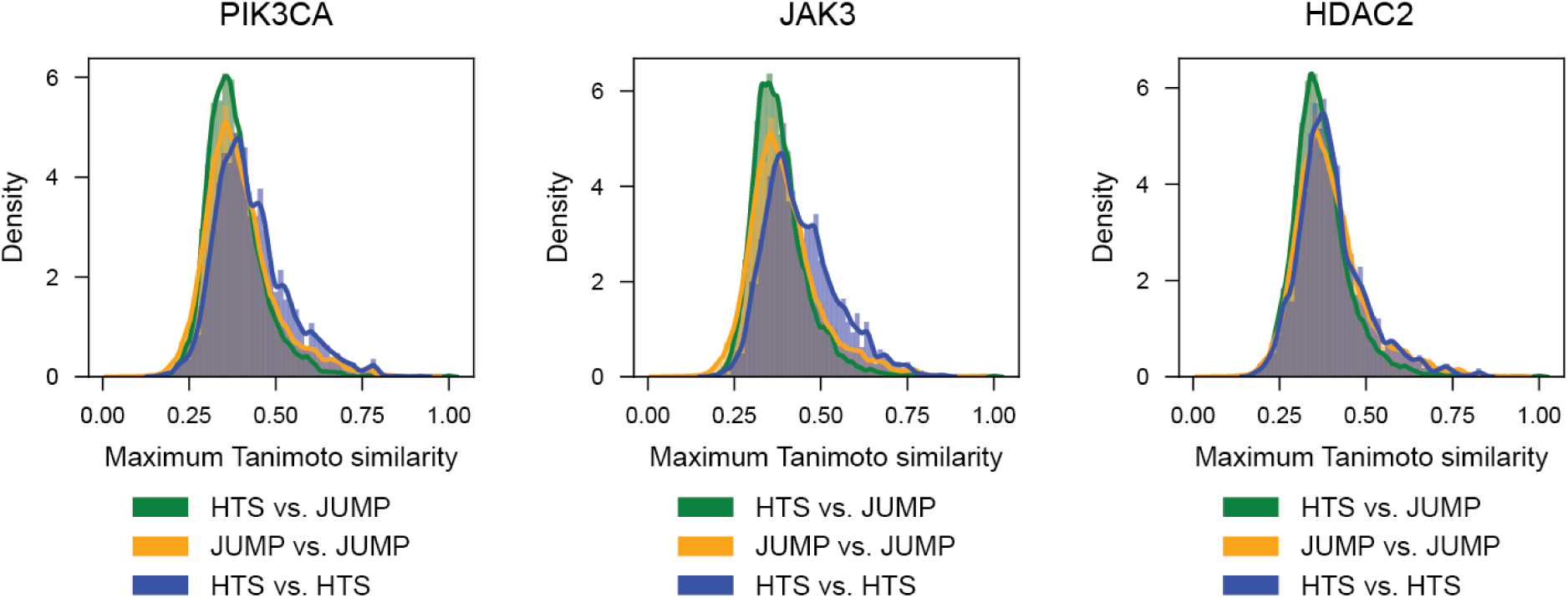
Distribution shift between JUMP and historical HTS datasets. Kernel density estimates (lines) and histograms (shaded areas) showing the distribution density (y axis) of the maximum Tanimoto similarities (x axis) between compounds in different datasets. For each compound in a given dataset, the maximum Tanimoto similarity is defined as the Tanimoto similarity of its most similar compound in the other dataset. Comparisons are shown for HTS versus JUMP, JUMP versus JUMP, and HTS versus HTS (colors) across three target datasets (panels), from left to right: PIK3CA, JAK3, and HDAC2.

**Fig. S9:**
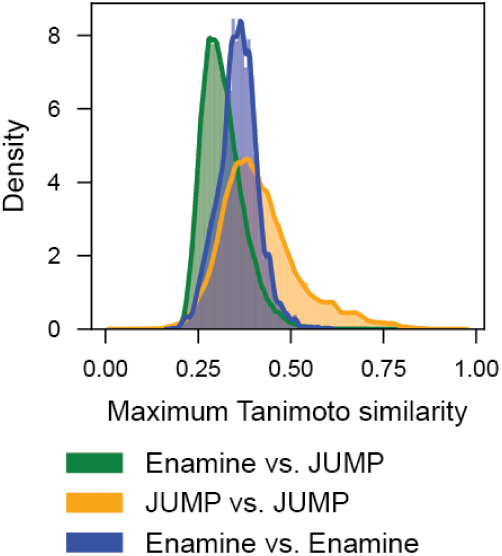
Distribution shift between JUMP and Enamine datasets. Kernel density estimates (lines) and histograms (shaded areas) showing the distribution density (y axis) of the maximum Tanimoto similarities (x axis) between pairs of compounds. Comparisons are shown for Enamine versus JUMP, JUMP versus JUMP, and Enamine versus Enamine (colors).

**Fig. S10:**
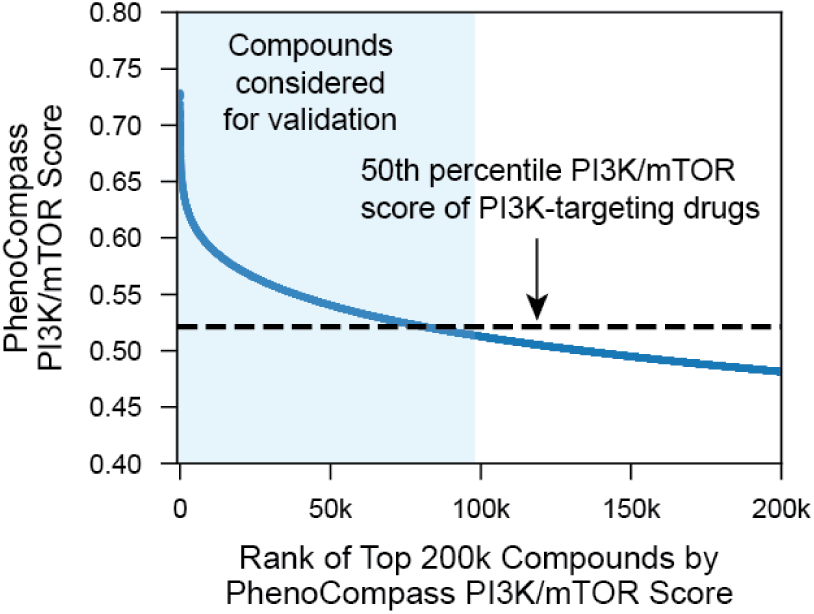
PhenoCompass PI3K/mTOR pathway scores for top Enamine REAL compounds. PhenoCompass PI3K/mTOR pathway score (y axis) plotted against the rank of the top 200,000 Enamine REAL compounds (x axis). The shaded region indicates the highest-scoring 100,000 compounds considered for experimental validation; the horizontal dashed line denotes the corresponding 50th percentile score of known PI3K-targeting drugs.

**Fig. S11:**
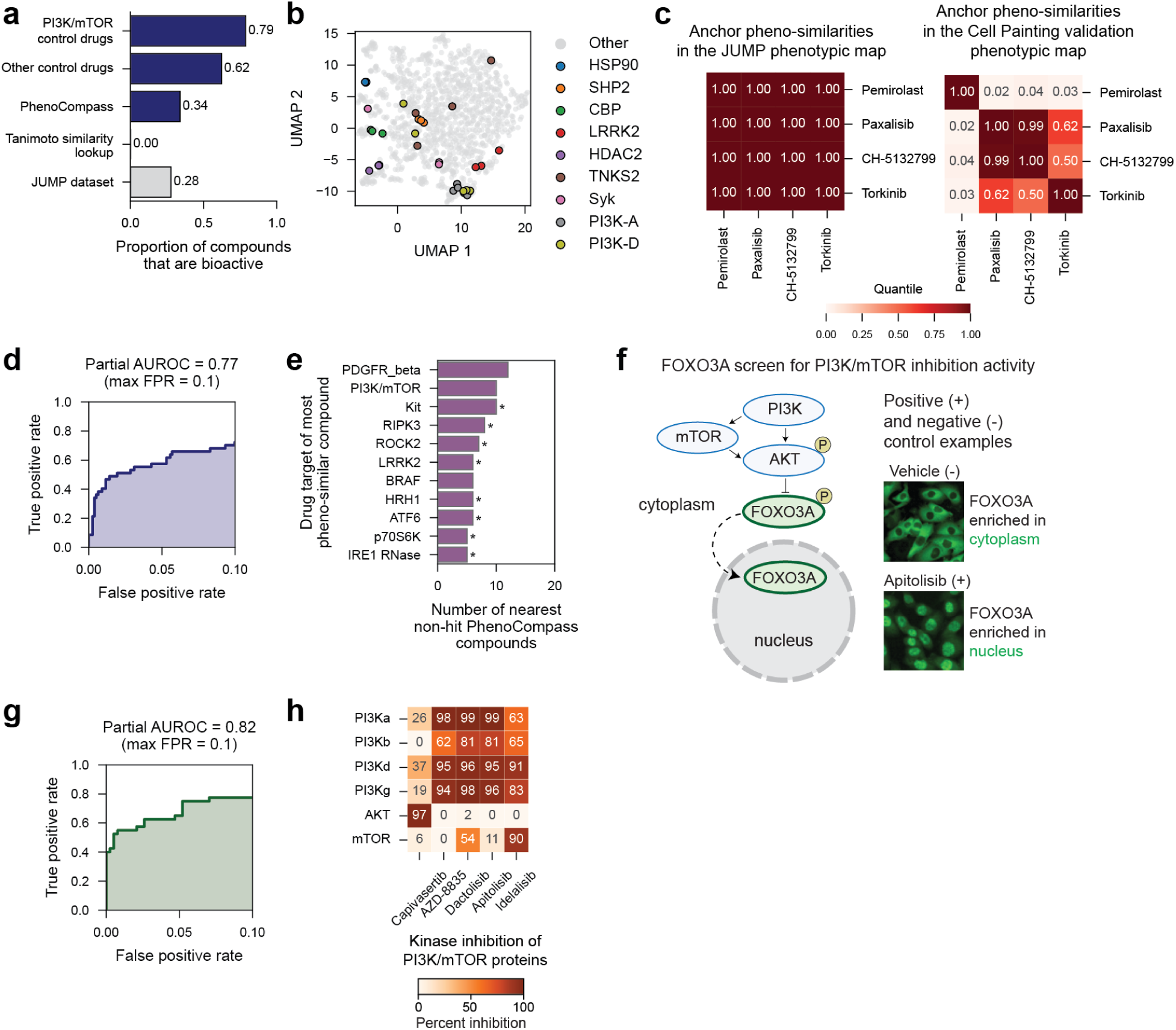
Details of the Cell Painting, FOXO3A translocation, and kinase inhibition validation experiments. **a**, Proportion of compounds that are bioactive (x axis) for different categories of compounds (y axis) screened in the Cell Painting validation data (blue bars) and the JUMP dataset (gray bars). **b**, UMAP visualization of mean-aggregated compound phenotypic profiles (dots) in the Cell Painting validation dataset. Colored dots correspond to top-9 MoAs by global biological recall (legend); gray dots correspond to PhenoCompass compounds and all control compounds that are not associated with the top-9 MoAs (**Methods**). **c**, Heatmaps of pheno-similarities (quantile-normalized cosine similarities) for anchor compounds present in both the JUMP dataset (left) and the Cell Painting validation dataset (right). Colors indicate similarity strength (color bar; low, blue; high, red). **d**, Accuracy of identifying known PI3K/mTOR-targeting compounds via Cell Painting score. Partial Receiver Operator Characteristic (ROC) curve (y axis, true positive rate versus x axis, false positive rate; max FPR = 0.1) using the mean cosine similarity to PI3K/mTOR anchor compounds in the Cell Painting validation map as a predictor of PI3K- or mTOR-targeting compounds against a background of annotated compounds targeting other genes. **e**, Potential off-target hits within PhenoCompass-predicted compounds. Number of nearest non-hit PhenoCompass compounds (x axis) that are pheno-similar to annotated control compounds, grouped by their target gene annotation (y axis). Asterisks (*) indicate statistically significant enrichment (*p* < 0. 05, hypergeometric test). **f**, Visualization of the PI3K/mTOR pathway and its relationship to FOXO3A translocation. Left: Diagram showing how inhibition of the PI3K/mTOR axis prevents FOXO3A phosphorylation, leading to its nuclear translocation. Right: representative images of negative (vehicle, cytoplasmic enrichment) and positive (Apitolisib, nuclear enrichment) control conditions. **g**, Accuracy of identifying known FOXO3A-active compounds via Cell Painting score. Partial ROC curve (y axis, true positive rate versus x axis, false positive rate) of the mean cosine similarity to PI3K/mTOR anchor compounds in the Cell Painting validation map as a predictor of FOXO3A active compounds. **h**, Heatmap of kinase inhibition percentages for specific kinase targets (rows) for positive control compounds (columns; color bar: low, white; high, orange).

**Fig. S12:**
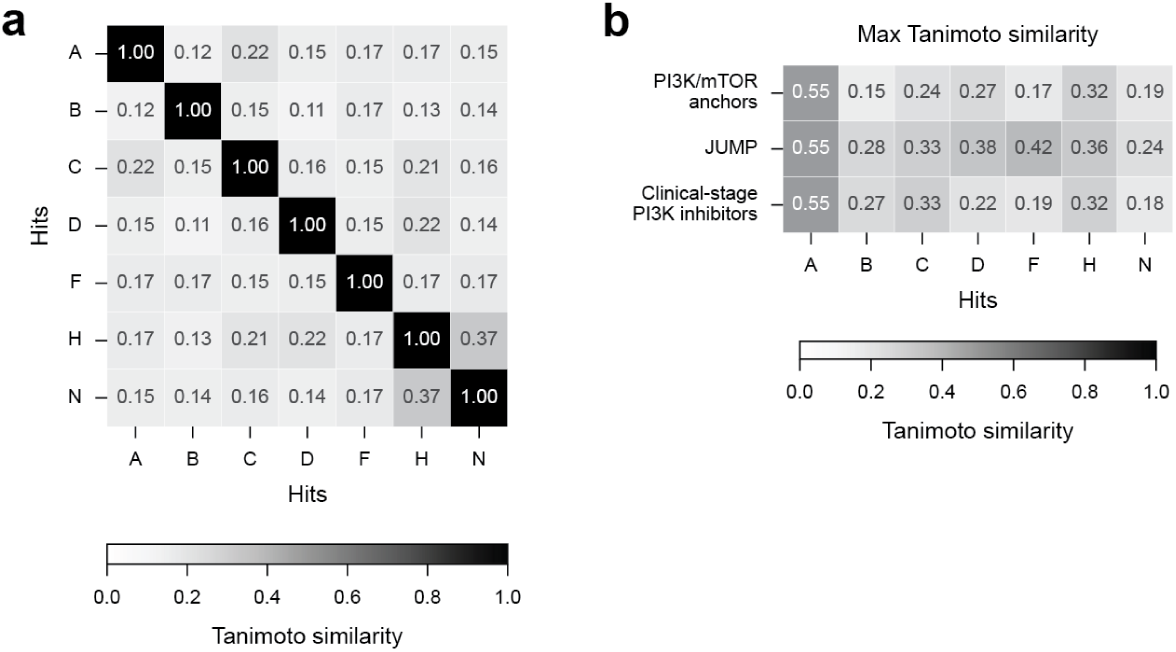
Structural similarities of kinase inhibition hit compounds. **a**, Heatmap of Tanimoto similarities (color bar; low, white; high, black) between all pairs of kinase inhibition hit compounds A–N (x and y axes). **b**, Maximum Tanimoto similarities (color bar; low, white; high, black) between the kinase inhibition hit compounds A–N (x axis) and each of the PI3K/mTOR anchor compounds, the JUMP dataset, and the set of clinical-stage PI3K inhibitors (y axis).

**Table S1:** Well position confoundedness of genetic perturbations. CRISPR knockouts and ORF overexpression perturbations are strongly confounded by well position, resulting in high replicate consistency and separability from random background (phenotype rate) irrespective of whether the target gene is expressed in U2OS cells.

|  | Compounds | CRISPR | ORF |
| --- | --- | --- | --- |
| Perturbations seen in single well position (%) | 0.0 | 94.9 | 99.4 |
| Phenotype rate<br>$p \leq 0.05$ nominal (%) | | | |
| All perturbations | 49.2 | 99.3 | 93.5 |
| Expressed genes |  | 99.4 | 93.6 |
| Unexpressed genes |  | 99.0 | 92.9 |
| Phenotype rate<br>$p \leq 0.01$ nominal (%) | | | |
| All perturbations | 36.7 | 97.8 | 85.8 |
| Expressed genes |  | 98.1 | 86.2 |
| Unexpressed genes |  | 96.8 | 84.5 |
| Phenotype rate<br>$p \leq 0.05$ FDR controlled (%) | | | |
| All perturbations | 42.0 | 98.6 | 89.9 |
| Expressed genes |  | 98.8 | 90.2 |
| Unexpressed genes |  | 98.1 | 88.7 |
| Phenotype rate<br>$p \leq 0.01$ FDR controlled (%) | | | |
| All perturbations | 27.6 | 95.8 | 76.7 |
| Expressed genes |  | 96.4 | 77.3 |
| Unexpressed genes |  | 94.0 | 75.2 |

**Table S2:**
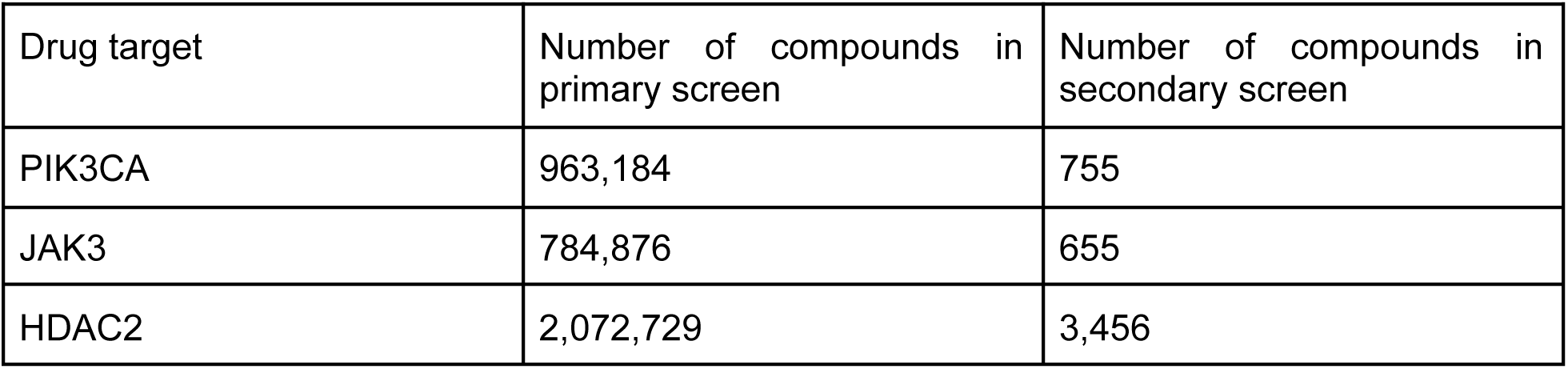
Number of compounds in each phase of the high-throughput binding affinity screens for each drug target.

**Table S3:** Hyperparameters for the PhenoCompass models.

| Model Component | Model Choice & Hyperparameters |
| --- | --- |
| **Networks** |  |
| GNEprop Compound Encoder Network | GNEprop: in-channels = 133, edge dimensions = 12, # of layers = 5, # of readout layers = 2, aggregation = global attention, dropout = 0 |
| Curated Features Compound Encoder Network | MLP: # of hidden layers = 1, # of hidden layer dimensions = 256, activation = ReLU |
| Compound Projection Network | MLP: # of hidden layers = 1, # of hidden layer dimensions = 256, activation = ReLU |
| Morphology Projection Network | MLP: # of hidden layers = 1, # of hidden layer dimensions = 256, activation = ReLU |
| Joint Projection Network | MLP: # of hidden layers = 1, # of hidden layer dimensions = 256, activation = ReLU |
| Common Encoder Network | MLP: # of hidden layers = 1, # of hidden layer dimensions = 256, activation = ReLU |
| **Contrastive Pre-Training Details** | Hyperparameters |
| GMC Objective + Consistency Regularization for PhenoCompass (GNEprop) Model | consistency loss weight $\alpha = 10$, temperature $\tau = 0.5$, mini-batch size = 256, optimizer = Adam, learning rate = 0.0001 |
| GMC Objective for PhenoCompass (Curated Features) Model | temperature $\tau = 0.2$, mini-batch size = 256, optimizer = Adam, learning rate = 0.0001 |

**Table S4:** Pre-processing step conditions included in the grid-search for constructing a phenotypic map from the Cell Painting validation data.

| Pre-processing step | Conditions |
| --- | --- |
| Feature normalization | Plate-wise z-scoring using (1) median as center + median absolute deviation (MAD) as dispersion, (2) median as center + interquartile range (IQR) dispersion |
| Dimensionality reduction | PCA with (1) 64, (2) 128, (3) 256 dimensions |
| Batch correction | (1) global whitening transformation, (2) plate-wise typical variation normalization |

**Data S1. (separate file)**

Source data underlying main and supplementary figures.

**Data S2. (separate file)**

Curated compound annotations (N = 1,239 compounds) across multiple resolutions (target gene, target gene family, target gene functional class) and ground-truth sources (OpenTargets, ChEMBL, ZINC15), used for pathway-anchor selection in the JUMP phenotypic map.

